# Circuit-specific dendritic development in the piriform cortex

**DOI:** 10.1101/868075

**Authors:** Laura Moreno-Velasquez, Hung Lo, Stephen Lenzi, Malte Kaehne, Jörg Breustedt, Dietmar Schmitz, Sten Rüdiger, Friedrich W. Johenning

## Abstract

Dendritic geometry is largely determined during postnatal development and has a substantial impact on neural function. In sensory processing, postnatal development of the dendritic tree is affected by two dominant circuit motifs, ascending sensory feedforward inputs and descending and local recurrent connections. In the three-layered anterior piriform cortex, neurons in the sublayers 2a and 2b display vertical segregation of these two circuit motifs. Here, we combined electrophysiology, detailed morphometry and Ca^2+^ imaging in acute mouse brain slices and modeling to study circuit specific aspects of dendritic development. We observed that determination of branching complexity, dendritic length increases and pruning occurred in distinct developmental phases. Layer 2a and layer 2b neurons displayed developmental phase specific differences between their apical and basal dendritic trees related to differences in circuit incorporation. We further identified functional candidate mechanisms for circuit-specific differences in postnatal dendritic growth in sublayers 2a and 2b at the meso- and microscale level. Already in the first postnatal week, functional connectivity of layer 2a and layer 2b neurons during early spontaneous network activity scales with differences in basal dendritic growth. During the early critical period of sensory plasticity in the piriform cortex, our data is consistent with a model that proposes a role for dendritic NMDA-spikes in selecting branches for survival during developmental pruning in apical dendrites. The different stages of the morphological and functional developmental pattern differences between layer 2a and layer 2b neurons demonstrate the complex interplay between dendritic development and circuit specificity.

**Significance Statement:** Sensory cortices are composed of ascending sensory circuits that relay sensory information from the periphery and recurrent intracortical circuits. Dendritic trees of neurons are shaped during development and determine which circuits contribute to the neuronal input space. To date, circuit-specific aspects of dendritic development and the underlying mechanisms are poorly understood. Here, we investigate dendritic development in layer 2 of the piriform cortex, a three-layered palaeocortex that displays a clear vertical segregation of sensory and recurrent circuits. Our results suggest that dendritic development occurs in distinct developmental phases with different circuit-specific properties. We further identify candidate mechanisms for neuronal activity patterns that could determine differences in circuit-specific dendritic development.

## Introduction

The complex geometry of neuronal dendritic trees in relation to their function is not yet fully understood. In sensory cortices, sensory input from the periphery is distributed to cortical neurons in an ascending sensory stream of input. Recurrent connectivity between cortical neurons constitutes the local and descending stream of input, which then transforms the sensory input into cortical output (Kanari et al., 2019; Srinivasan and Stevens, 2018). Developmental growth patterns of dendritic structures are an important determinant of a neuron’s function within the different circuits constituting its synaptic input space (Lanoue and Cooper, 2018). This brings up the question of how dendritic morphology develops in relation to the two different glutamatergic circuit elements in sensory information processing, ascending sensory input and recurrent connectivity.

We investigated the palaeocortical three-layered anterior piriform or primary olfactory cortex (aPCx), which shares structural and functional similarities with the reptilian dorsal cortex (Fournier et al., 2015). The aPCx is the largest cortical region receiving olfactory sensory inputs. Peripheral odor information from nasal olfactory sensory neurons converges onto the aPCx via the olfactory bulb. Functionally, the aPCx synthesizes the segregated peripheral input into odor objects and identifies them (Wilson and Sullivan, 2011). Unlike topographically organized neocortical sensory systems, afferent sensory and recurrent input streams to the aPCx lack any apparent spatial structure and are therefore non-topographical (Srinivasan and Stevens, 2018). Layer 2 is the main cellular layer of the olfactory cortex (Bekkers and Suzuki, 2013). Based on the distribution of genetic markers and somatic morphology, layer 2 can be divided into layer 2a (superficial third) and layer 2b (deeper two-thirds)(Bolding et al., 2019; Choy et al., 2017; Diodato et al., 2016; Martin-Lopez et al., 2017). Layer 2a predominantly contains superficial so-called semilunar cells (layer 2a neurons). Layer 2b harbors pyramidal cells and semilunar-pyramidal transition cells (layer 2b neurons) (Choy et al., 2017; Suzuki and Bekkers, 2011). Neurons in the two sublayers display differences in functional circuit incorporation. Layer 2a neurons predominantly sample converging sensory input and distribute it unidirectionally to the layer 2b and 3 neurons. Layer 2b neurons receive sensory input and, in addition, are incorporated in a rich recurrent network (Choy et al., 2017; Hagiwara et al., 2012; Suzuki and Bekkers, 2011; Wiegand et al., 2011). Recently, it has been demonstrated in vivo that these two neuron types play different roles in reading out converging sensory input (layer 2a neurons) and performing pattern storage and completion via recurrent circuits (layer 2b neurons) (Bolding et al., 2019).

This vertical organization of input space of layer 2 neurons extends to the dendritic tree, where sensory and recurrent functional domains are spatially segregated. In the apical dendrites of all neurons in layer 2, the majority of sensory input projects to the superficial layer 1a. Layer 1a can be clearly distinguished from layer 1b, which, together with inputs in layer 2 and 3, samples recurrent inputs. Basal dendrites exclusively sample recurrent inputs (Kevin M. Franks and Isaacson, 2005; Johenning et al., 2009). In aPCx, we therefore observe a clear vertical segregation of functionally distinct cell types and of different functional dendritic domains. This feature of aPCx makes layer 2 of the aPCx an ideal model for the differential analysis of dendritic growth patterns related to sensory input and recurrent connectivity.

Here, we studied developmental dendritic growth in layer 2a and layer 2b neurons in acute brain slices of the aPCx. We applied electrophysiology, detailed morphometry of 3D-reconstructed neurons, Ca^2+^ imaging and computational modelling. We identified distinct phases of dendritic development with cell-type specific differences of dendritic growth and pruning patterns. We related the different developmental patterns described here at the morphological level to physiological differences at the micro- and mesoscale level. This enabled us to identify candidate mechanisms that may drive circuit-specific dendritic development in a non-topographic sensory system.

## Materials and Methods

### Slice preparation

Acute brain slices were prepared from C57Bl6N mice of either sex except for population Ca^2+^ imaging experiments with GCaMP (Fig. 4), where Ai95-NexCre mice were used. In experiments for Figs. 1-5, the horizontal slicing orientation was chosen to preserve rostrocaudal association fibers (Demir et al., 2001). For layer 1a dendritic spike measurements in Fig. 6, we used coronal slices. All animal procedures were performed in accordance with the [Author University] animal care committee’s regulation. For morphological reconstruction, acute brain slices were prepared at 4 age intervals: p1-2, p6-8, p12-14 and p30-40 (>p30). Electrophysiological Characterization was limited to the 2 age intervals p12-14 and p30-40. For measurements of NMDA-spikes, coronal slices were prepared at p14-21. Brains from p30-40 mice and from mice used for dendritic spike measurements were prepared in ice-cold artificial cerebrospinal fluid (ACSF; pH 7.4) containing (in mM): 87 NaCl, 26 NaHCO3, 10 Glucose, 2.5 KCl, 3 MgCl2, 1.25 NaH2PO4, 0.5 CaCl2 and 50 sucrose. Slices were cut at 400 µm thickness, and incubated at 35°C for 30 min. The slices were then transferred to standard ACSF containing (in mM): 119 NaCl, 26 NaHCO3, 10 Glucose, 2.5 KCl, 2.5 CaCl2, 1.3 MgCl2, and 1 NaH2PO4. Slices from other age groups were cut in ice-cold standard ACSF and incubated for 30 min in standard ACSF at 35°C. The slices were then stored in standard ACSF at room temperature in a submerged chamber for 0.5–6 h before being transferred to the recording chamber. For dendritic spike measurements in Fig. 6, 1 µM of Gabazine was added to the recording solution. Experiments requiring spontaneous network activity were prepared in ice-cold ACSF containing (in mM): 125 NaCL, 25 NaCHCO3, 10 Glucose, 4 KCL, 1.25 NaH2PO4, 2 CaCl2, 1 MgCl2. Slices were incubated at 35°C for 30 min and stored at room temperature in a submerged chamber for 0.5–7 h. All recordings were performed at near-physiological temperature (32-34 °C).

**Figure 1.**
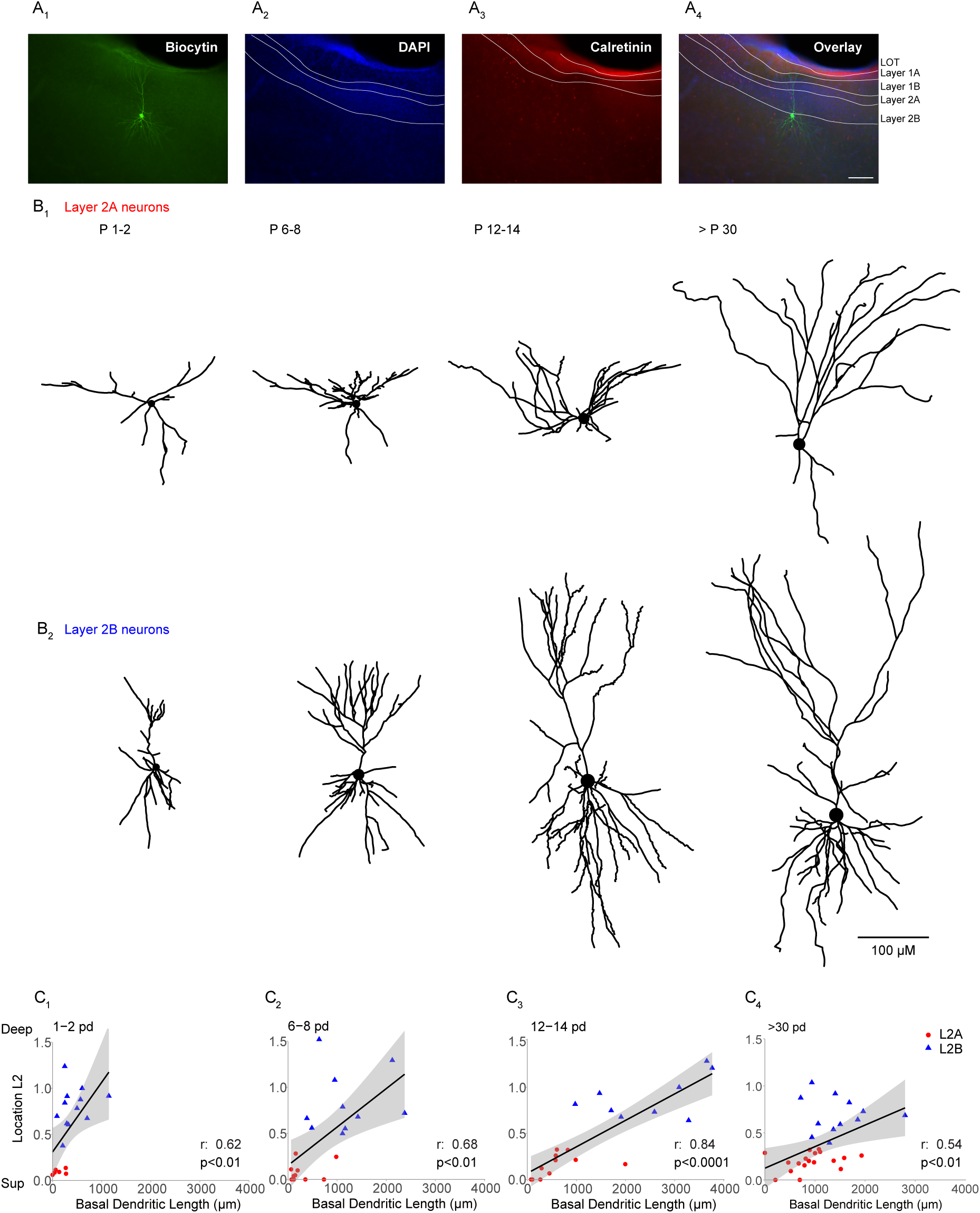
Localization and differentiation of the two principal neuron types in Layer 2 of the aPCx. (A) Slice containing one recorded neuron filled with biocytin (A1) and additionally, post hoc labelled with DAPI (A2) and calretinin (A3). The overlay (A4) shows the recorded neuron located in the layer 2b/layer 3 transition zone of the aPCx. (B) 3D Morphological reconstructions of different layer 2a (B1) and layer 2b (B2) neurons at four time windows: Right after birth (p1-2), at the end of the first postnatal week (p6-8), at the end of the second postnatal week (p12-14) and after the fifth postnatal week (>p30). Scalebar 100 µm. (C) Correlation between the total basal dendritic length and the vertical position of the cells in layer 2 are shown at the same four time windows. Spearman r and p-values are shown for each time group.

#### Electrophysiology

Whole-cell current clamp experiments were performed at near physiological temperature (32-34 °C) using an Axon Multiclamp 700B amplifier (Molecular Devices, Sunnydale, CA, US). For morphological reconstruction and characterization, signals were low pass filtered at 2 kHz and digitized at a sampling rate of 20 kHz (BNC-2090, National Instruments Corporation, Austin, Tx, US). Pipettes (3-6 MΩ) were filled with an intracellular solution containing (in mM): 135 K-gluconate, 6 KCl, 10 HEPES, 0.2 EGTA, 2 MgCl2, 2 Na-ATP, 0.5 Na-GTP, 5 phosphocreatine Na (pH: 7.3) and Biocytine (0.20%). Liquid junction potential (LJP) was not corrected. Bridge balance compensation was applied in current clamp. Cells were discarded if the resting membrane potential was above -60 mV or the series resistance exceeded 30 MΩ. For dendritic spike recordings, signals were low pass filtered at 8 kHz and digitized at a sampling rate of 20 kHz. Pipettes (3-6 MΩ) were filled with an intracellular solution containing (in mM): 130 K-gluconate, 20 KCl, 10 HEPES, 4 MgATP, 0.3 NaGTP and 10 phosphocreatine (pH:7.3, adjusted with KOH), 30 µM Alexa 594 and 500 µM fluo-5F. Experiments were conducted without exceeding -200 pA at resting membrane potential (Layer 2a neurons were held at -60 mV, and layer 2b neurons were held at -70 mV). Series resistance was below 30 MΩ. After dye-filling the patched neuron for 10 mins, we placed the theta glass stimulation electrode close to the distal dendrite in layer 1a of the piriform cortex. Stimulation protocol was set to 3 pulses at 50 Hz with 10 µA steps (with one exception in layer 2b neuron, which was with 20 µA steps.).

#### Electrophysiological Analysis

Analysis was performed using custom-written routines in Python. Resting membrane potential (Vm) was taken as the mean value of the baseline before current injections were performed. For characterization, neurons were held at -60 mV. Input resistance (IR), membrane time constant (Tau) and membrane capacitance (Cm) were calculated from the voltage response to an 80 pA hyperpolarizing current step. Action potential (AP) threshold was defined as the membrane potential at the point where the slope (dV/dt) reached 1 % of its maximum. The fast after-hyperpolarisation (fAHP) was defined as the difference between AP threshold and the minimum voltage seen immediately after the AP peak (within 5 ms). Finally, the instant firing frequency was defined as the frequency between the first and second AP. For comparability, these values were extracted from the first 600 ms current injection step that elicited at least 9 APs. When analyzing the integrative behavior of apical dendrites in Fig. 6, we analyzed the changes in EPSP size upon linear increase of stimulation intensity. To quantify EPSP size, we measured the amplitude and area under curve of a 60 ms time window following the 3rd pulse compared to baseline (50 ms period before stimulus). Effects of APV were quantified for the largest response that did not yet evoke an AP.

#### Immunohistochemistry

Slices with biocytin-filled cells were stored in 4 % paraformaldehyde (PFA) overnight. The following day, slices were washed 3 times (10 mins each) in PBS and incubated in a blocking solution composed of 5 % normal goat serum (NGS, Biozol), 1 % Triton-X (Sigma) and PBS, for 3 hours at room temperature with gentle agitation. Primary antibodies were diluted in blocking solution (2.5 % NGS, 1 % Triton-X, PBS) and slices were incubated for 72 hours at 4 °C. Biocytin-filled neurons were labeled with a streptavidin marker conjugated to Alexa Fluor (AF) 488 (Invitrogen, S-32354; 1:500 dilution). Additionally, the LOT and mitral cell axons in layer 1a were labeled with calretinin (anti-mouse; Millipore, MAB1568; 1:1000 dilution or anti-rabbit; SWANT, 7697; 1:4000 dilution) and interneurons with GAD 67 (anti-mouse; Millipore, MAB 5406; 1:500 dilution), GAD 65/67 (anti-rabbit; Chemicon, AB 11070; 1:500 dilution) or Gephyrin (anti-mouse; SYSY 147 111; 1:500 dilution).

Following this, slices were washed 2 times (10 mins each) with PBS and secondary antibodies (goat anti-rabbit AF 555, goat anti-rabbit AF 647, goat anti-mouse AF 555, goat anti-mouse AF 647; Invitrogen; 1:500 dilution in 0.5% Triton, PBS) were applied for 3 hours at room temperature. Finally, slices were washed 3 times (10 mins each) in PBS and mounted on glass slides in mounting medium Fluoroshield with 4’, 6-diamidino-2-phenylindole (DAPI; Sigma).

### Reconstructions and Morphological Analysis

Mounted slices were visualized on a fluorescent microscope (10x objective, 0.3N.A.; Leica) in order to identify and select the biocytin-filled neurons located in the aPCx for further reconstruction. Only neurons that displayed homogenous filling with Biocytin and lacked obvious amputation of the dendritic tree by slicing were analyzed. Therefore, not all neurons chosen for electrophysiology were also chosen for morphological reconstruction and vice versa. Selected slices were then imaged on an upright Leica TCS SP5 confocal microscope (Leica Microsystems) through a 20x immersion objective (0.7 N.A.; Leica) with 405 nm (diode), 488 nm (Argon laser), 568 nm (solid state) and 633 nm (Helium, Neon) laser lines. For biocytin-filled neurons, the perisomatic field of view was further imaged through a 63x immersion objective (1.4 N.A.; Leica) to validate the spine density. Cells were selected and classified according to their position in layer 2 of the anterior piriform cortex using FIJI (https://imagej.nih.gov/ij/). The position in layer 2 was defined as the smallest distance from the soma to the border between layer 1b and layer 2a, normalized to the total width of layer 2 for each neuron. The border between layer 1a and 1b was traced to later classify the apical dendrites according to their synaptic inputs. Neuronal morphologies were then reconstructed with neuTube software (Feng et al. 2015) and exported as SWC files. Morphometric parameters were extracted with L-measure software (Scorcioni et al., 2008) and analysed with R studio and Python using btmorph v2 (Torben-Nielsen, 2014) and scipy packages.

#### Ca^2+^ Imaging

For population Ca^2+^ imaging of neonatal spontaneous synchronous network events (Fig. 4), we used the genetically encoded Ca^2+^ indicator (GECI) GCaMP6F. NEX-Cre mice (Goebbels et al., 2006) were crossed with Ai95 animals (https://www.jax.org/strain/024105 (Madisen et al., 2015)) for constitutive GCaMP6F expression in excitatory cells only.

For experiments involving spontaneous network activity, Ca^2+^ imaging was performed using a Yokogawa CSU-22 spinning disc microscope at 5000rpm. The spinning disc confocal permitted the generation of a large field of view time series at a high acquisition rate. A 488 nm Laser was focused onto the field of view using a 40x objective. Emission light was filtered using a 515±15nm bandpass filter. Fluorescence was detected using an Andor Ixon DU-897D back-illuminated CCD, with a pixel size of 16µm. Andor iQ software was used for data acquisition. Population Ca^2+^ imaging was performed at 10 Hz when single cells were measured. For measuring of larger ROIs incorporating layer 2, 10 Hz data was pooled with a dataset acquired at 40 Hz.

For analysing dendritic spikes using 2P-imaging, 30 µM Alexa-594 and 500 µM Fluo-5F were added to the intracellular solution. A Femto 2D two-photon laser scanning system (Femtonics Ltd., Budapest, Hungary) equipped with a femtosecond pulsed Ti:Sapphire laser tuned to λ=805 nm (Cameleon, Coherent, Santa Clara, CA, US) controlled by the Matlab-based MES software package (Femtonics Ltd., Budapest, Hungary) was used. Fluorescence was detected in epifluorescence mode with a water immersion objective (LUMPLFL 60x/1.0 NA, Olympus, Hamburg, Germany). Transfluorescence and transmitted infra-red light were detected using an oil immersion condenser (Olympus). The average scanning speed was 300 Hz and the intermediate sections were jumped over within 60 µs using a spline interpolated path. Dendritic Ca^2+^ transients were measured every 30 s.

#### Imaging Analysis

For population Ca^2+^ imaging (Fig. 4), FOVs with at least 5 minutes of recordings were included in the analysis. Videos were motion corrected using Suite2p (Pachitariu et al., 2016) and analysed using custom python code and SamuROI (Rueckl et al., 2017). If motion artefacts were too great to be corrected, recordings were not included in the subsequent analysis. Each pixel of the raw data was normalised using the 6-sample window with the lowest standard deviation. Traces were extracted from each ROI and event detection was carried out. Events were detected as increases in ΔF/F greater than 2.5 standard deviations from baseline with a peak width of at least 2 consecutive samples. The results were manually curated with the user free to exclude events based on the inter-event-interval, amplitude, signal to noise ratio and peak width. Incomplete events at the start or end of each recording were excluded from analysis. The rates of false positives and negatives were 4.5% and 5.1% respectively calculated from a random subset of the data (100 cells, 3 mice).

The upper and lower boundaries of layer 2 were manually defined based on cell density. For detection of global events, we measured the average change in fluorescence for all pixels of layer 2 piriform cortex, including the neuropil, using a rectangular ROI defined by the upper and lower boundaries of layer 2.

For single cell analysis we used a semi-automated method based on image segmentation with Ilastik (Sommer et al., 2011). This was required because a large number of cells were inactive, or closely packed and/or synchronous in their activity. Ilastik was trained to segment z-projection sum images of a subset of FOVs to produce a 5-label image (nuclei, somata, bright debris, dark debris and background). Cells were detected using the nuclei label with false positives manually rejected. Using these cell locations, the somata image was divided into territories using watershed segmentation and only the nearest pixels to each nucleus were included. ROIs with fewer than 70 pixels were rejected. We calculated the ΔF/F for each ROI and subtracted an estimate of the local neuropil contribution using an equal number of randomly selected non-cell pixels within a fixed radius of 70 pixels.

#### Modelling

For the model in Fig. 7, we assumed that the density of synapses on the branches of a dendrite is constant, i.e. depends linearly on the length of a branch. We assumed that upon odor exposure a maximum of 70 active synapses can arrive at a single neuronal dendrite. Following recent experimental measurements (Srinivasan and Stevens, 2018), we considered 3700 glomeruli, 9.7×10^7^ synapses between all glomeruli and all layer 2 neurons and a total of 41000 layer 2 neurons. Hence, we dealt with an average 9.7×10^7^ / 41000 = 2366 synapses between all glomeruli and one neuron. Thus for 70 synapses to be activated upon odor exposure, we assumed that 109 glomeruli are activated per odor (=3700×70/2366).

We characterized the morphology of a neuron through its mean branch length (BL), which we found to range from 40 to 110 µm (mean is 72 µm), cf. Results. The total dendritic branch length on average is approximately 1800 µm.

For simplicity, we approximated the length of any branch by the mean values of the respective neurons. Accordingly, dendrites could have a maximum of 1800/40 = 45 (short-branched neuron) and a minimum of 1800/110 = 16 branches (long-branched neuron).

We distinguished the two cases of clustered (n_clus_) and distributed (n_dist_) stimulation. We assumed that a neuron fires if it is exposed to more than n_dist_=40 active input synapses or n_clus_=10 active inputs arriving on the same branch. The excitation behavior is schematically illustrated in Figure 7A, showing a simplified version of the non-linear dendritic integration scheme proposed by Poirazi et al. (Poirazi et al., 2003). Specifically, we considered every neuron as a two-layered network that may or may not produce a network response (express a somatic action potential) to a presented stimulus set. This response is triggered in cases when the distributed input reaches a certain threshold number (reflecting a number of active input synapses), which can be understood as a linear integration scheme of the neuron. In addition, the network may produce a response to a dendritic spike. In our model, dendritic spikes led to the non-linear integration of synaptic input, which is mimicked through the activation gates on every branch of the neuron (first layer of the network). In Fig. 7A, this first layer of the network is shown as blue circles. The green circle represents the soma. The magnification insets illustrate the stimulus response relationship of the separate branches and the soma, respectively. The model was constructed such that a somatic response is expressed, if the distributed stimulation crosses the threshold value n_dist_, or if one of the branches expresses a dendritic spike, which relates to the number of active synapses on the branch crossing the threshold number of clustered stimulation n_dist_.

Using this described model, we investigated the additional response probability that is introduced through considering clustered stimulation. We supposed the number of odor-activated synapses (which we refer to as) connected to a given neuron to be random. The probability of finding a neuron that is connected to or more odor activated synapses can then be approximated as

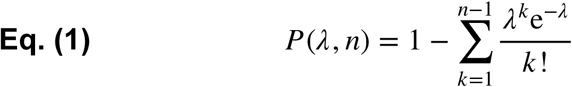

Eq. (1) allowed us to map the response probability of a neuron to a presented stimulus. In the case of clustered stimulation, this picture is slightly different: Instead of the mean number of synapses per neuron, the mean number of synapses per branch is the relevant quantity. The mean number of branches (NB) is approximately the total dendritic branch length (TDBL) divided by the mean branch length (BL). We took the distribution of BL into account as we reasoned that longer branches optimize the input-output relationship in case of clustered stimulation. At P12-14, the length of the dendritic branches ranged from 40 µm to 110 µm. The response probability for clustered stimulation is modeled as:

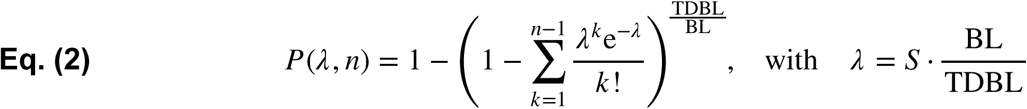

It is important to note that under such a scheme, stimulation of any dendritic branch can be sufficient to excite the neuron.

## Code accessibility

The code underlying the calculations and plot in Fig. 7 is available as Extended Data 1 and via github (https://github.com/mkahne/DendriticBranches). The code was executed using Python 3.7.3 on a macbook pro running on MacOs 10.15.4.

## Statistics

Data were first tested for normality. Statistical tests were performed as indicated using GraphPad prism, the scipy library and the DABEST package in Python and R (Ho et al., 2019). We used the paired (normally distributed direct comparisons) and unpaired t-test (normally distributed single comparisons), Wilcoxon test (not normally distributed direct comparisons), Mann-Whitney test (not normally distributed single comparisons), one-way ANOVA with Holm-Sidak’s multiple comparisons test (normally distributed multiple comparisons) or Kruskal–Wallis test with Dunn’s multiple comparison as a post hoc test (not normally distributed multiple comparisons) as indicated in the text. Extended Data Table 1-2 contains all applied tests and the exact p values. Additionally, the spearman correlation test was used to measure the association between cell position and branch-length in Fig. 1C, statistical details can be found in Extended Data Table 1-2. Numerical values are given as mean and SEM unless otherwise stated. To facilitate the interpretation of our results and the narrative flow of the paper, we followed the convention of defining p < 0.05 as significant in the text. However, in order to facilitate the realistic evaluation of our data and its interpretation, we omitted significance stars from the majority of the plots. Wherever possible, we used estimation-based statistics with mean-difference plots instead (Ho et al., 2019).

## Results

### Layer identification in postnatal development

In acute horizontal mouse brain slices, we performed whole cell patch clamp recordings of randomly sampled excitatory neurons over the whole vertical extent of layer 2 including the layer 2/3 transition zone. Excitatory neurons were distinguished from interneurons by at least one of the 3 criteria: firing profile (Suzuki and Bekkers, 2010), morphology and a negative post-hoc staining for interneuron markers. During patching, neurons were filled with biocytin for later morphological reconstructions (Fig. 1A1).

For analysis, the extent of layer 2 was delineated using a DAPI stain (Fig. 1A2). Layer 2 was divided into layer 2a (upper third) and layer 2b (deep two-thirds and layer 2/3 transition zone) (Choy et al., 2017; Martin-Lopez et al., 2017). We chose this division into sublayers for categorization although there is a postulated gradient of the electrophysiological and morphological differences between deep and superficial neurons (Suzuki and Bekkers, 2011; Wiegand et al., 2011). Our rationale behind this hard segregation is the clear distribution of genetic markers in layer 2 and the necessity to find a criterion applicable to all age groups studied. All published genetic markers for superficial or layer 2a neurons seemed to display a clear and selective expression profile in the upper third of layer 2 (Bolding et al., 2019; Choy et al., 2017; Diodato et al., 2016). In addition, functional analysis of neurons expressing genetic markers of layer 2a neurons display reduced recurrent circuit incorporation (Bolding et al., 2019; Choy et al., 2017).

We confirmed the location of the aPCx by staining the LOT and layer 1a fibers with calretinin (Sarma et al., 2011) (Fig. 1A3). The calretinin staining also permitted clear delineation of dendritic segments terminating in layer 1a (Fig, 1A4, see Fig. 5). 3D reconstructions using neutube (Feng et al., 2015) of layer 2a (Fig.1B1; n=46/25 neurons/mice) and layer 2b (Fig.1B2; n=43/27 neurons/mice) neurons were analyzed at four time windows: Right after birth (p1-2, layer 2a: n=6/4; layer 2b: n=12/6; neurons/ mice), at the end of the first postnatal week (p6-8, layer 2a: n=10/5; layer 2b: n=10/7; neurons/ mice), during the critical period of heightened sensory synaptic and structural plasticity (Franks and Isaacson 2005; Poo and Isaacson 2007; p12-14, layer 2a: n=11/8; layer 2b: n=9/8; neurons/mice) and after the critical period (>p30, layer 2a: n=19/6; layer 2b: n=12/6; neurons/mice; Fig. 1B).

### Neurons in sublayers 2a and 2b are distinct

Electrophysiological characterization was performed for a subset of neurons at p12-14 (layer 2a: n=10/6; layer 2b: n=8/7; (neurons, mice)) and at >p30 (layer 2a: n=14/7; layer 2b: n=9/6; (neurons, mice)). When sampling the whole extent of layer 2a and layer 2b, we did not find statistically significant electrophysiological differences between neurons in the layers 2a and 2b at both ages (Table 1,Extended Data Table 1-1, Extended Data Table 1-2; see discussion).

In addition to previously reported electrophysiological differences, the less complex basal dendritic tree of superficial layer 2a cells compared to deeper neurons in layer 2b is a prominent distinctive feature (Bekkers and Suzuki 2013). As basal dendritic length most likely scales with local recurrent wiring (Haberly, 1985), this is a good indicator of a layer 2 neuron’s recurrent circuit incorporation, which is central to this study. After the first postnatal week, we saw differences in the architecture of the basal dendritic tree between layer 2a neurons and layer 2b neurons. When plotting the normalized cell depth in layer 2 against the total dendritic branch length of each neuron’s basal tree, we observed a stable superficial to deep gradient of basal dendritic tree length and complexity over postnatal development (Fig. 1C, Extended Data Fig. 1-1; p1-2 (r=0.62, p<0.01), p6-8 (r=0.68, p<0.01), p12-14 (r=0.84, p<0.0001) and >p30 (r=0.54, p<0.01); spearman correlation.). When using our positional grouping approach, the number of basal branches and the total basal dendritic branch length were significantly smaller in layer 2a than in layer 2b neurons starting in postnatal week 1 (see also Fig. 3A; number of branches: layer 2a vs layer 2b: p6-8: p<0.001, p12-14: p<0.0001, >p30: p<0.01; total basal dendritic length: layer 2a vs layer 2b: p6-8: p<0.01, p12-14: p<0.0001, >p30: p<0.05; ANOVA with Holm-Sidak’s multiple comparisons test). We conclude that the morphological parameter basal dendritic length justifies the distinction between a superficial (layer 2a) and deep (layer 2b) population of aPCx layer 2 neurons in our dataset.

**Table 1.** Intrinsic electrical properties of layer 2a and layer 2b neurons at p12-14 and > p30. SL refers to layer 2a neurons, SP to layer 2b neurons.

**SL refers to layer 2a neurons, SP to layer 2b neurons.**
|  | 12-14 pd |  | >30 pd |  |
| --- | --- | --- | --- | --- |
|  | SL | SP | SL | SP |
| **V<sub>m</sub> (mV)** | -75.32 ± 1.96 | -73.48 ± 2.86 | -71.88 ± 1.89 | -72.23 ± 2.01 |
| **R<sub>in</sub> (MΩ)** | 233.35 ± 40.24 | 147.52 ± 27.07 | 168.71 ± 22.65 | 235.29 ± 51.62 |
| **Tau (ms)** | 25.65 ± 2.94 | 22.98 ± 4.24 | 20.79 ± 1.98 | 22.53 ± 1.48 |
| **C<sub>m</sub> (pF)** | 123.17 ± 11.86 | 156.30 ± 10.44 | 135.89 ± 9.66 | 114.52 ± 10.43 |
| **Threshold (mV)** | -35.97 ± 1.50 | -39.37 ± 1.17 | -40.28 ± 1.52 | -40.35 ± 1.07 |
| **FAHP (mV)** | 7.75 ± 1.30 | 5.65 ± 1.51 | 10.83 ± 1.59 | 7.96 ± 0.77 |
| **Instant Firing Freq (Hz)** | 37.99 (22.69-108.13) | 34.81 (29.90-104.40) | 44.87 (30.83-197.12) | 68.78 (29.42-88.44) |

median (IQR)

### Distinct growth phases in apical and basal dendrites

We chose a set of morphometric parameters to describe the growth of the apical (Fig. 2) and basal (Fig. 3) dendritic tree: number of branches, total dendritic branch length, number of stems, average individual branch length and branch density as a function of distance from the soma. Using these parameters, we defined three distinct developmental phases. In apical dendrites of both layer 2a and layer 2b neurons, we observed the largest fractional increase in branch number in the first postnatal week (layer 2a: 75% of total increase in branch number, layer 2b: 90% of total increase in branch number, Fig. 2A2). For layer 2b neurons, we observed a significant increase in apical branch number between p0-2 and p6-8 (Fig. 2A2; layer 2b neurons: p1-2 vs p6-8: p< 0.001, p12-14 vs >p30: p<0.05; ANOVA with Holm-Sidak’s multiple comparisons test). Layer 2a neurons displayed a significant addition of proximal stems in the first postnatal week only (Fig. 2A4; layer 2a neurons: p1-2 vs p6-8: p< 0.01, p12-14 vs >p30: p<0.01; ANOVA with Holm-Sidak’s multiple comparisons test). Layer 2b neuron basal branches displayed a similar developmental pattern (Fig. 3A2). Here, we observed a statistically significant increase in branch number in the first postnatal week (layer 2b: 64% of total increase in branch number, Fig. 3A2). In the shorter and less complex basal tree of layer 2a neurons (see above), significant branch addition was only evident when compared over the whole developmental period observed (Fig. 3A2; layer 2b neurons: p1-2 vs p6-8: p<0.01, p12-14 vs >p30: p< 0.05; layer 2a neurons: p1-2 vs p30: p<0.05; ANOVA with Holm-Sidak’s multiple comparisons test).

After this initial determination of branch complexity by branch addition (developmental phase 1), dendrites grew by branch elongation (developmental phase 2). In layer 2a neurons, we observed statistically significant increases of the total apical dendritic branch length in the second postnatal week and between weeks 2 and 5 (Fig. 2A3). In layer 2b apical dendrites, the total dendritic branch length increased significantly in the first and second postnatal weeks (Fig. 2A3; (layer 2a neurons: p6-8 vs p12-14: p<0.05, p12-14 vs >p30: p<0.05; layer 2b neurons: p1-2 vs p6-8: p<0.05, p6-8 vs p12-14: p<0.01; ANOVA with Holm-Sidak’s multiple comparisons test). In basal dendrites of layer 2b neurons, we observed increases in total dendritic branch length in developmental phase two until p12-14 (Fig. 3A3). In the shorter and less complex basal tree of layer 2a neurons, significant length growth was again only evident when comparing over the whole developmental period observed (Fig. 3A3; layer 2a neurons: p1-2 vs >p30: p<0.05; layer 2b neurons: p0-2 vs p6-8: p<0.05, p6-8 vs p12-14: p<0.001, p12-14 vs >p30: p<0.001; ANOVA with Holm-Sidak’s multiple comparisons test).

Increases in the total dendritic branch length are a combined effect of branch addition and elongation of individual branch segments. The dichotomy between branch addition in developmental phase one and the length growth of existing branches in developmental phase 2 became apparent when examining the average branch length per neuron (Figs. 2B and 3B). In layer 2a and layer 2b neuron apical and basal branches, the average branch length per neuron did not increase in the first postnatal week. In the second postnatal week and between weeks 2 and 5, layer 2a and layer 2b neurons both displayed significant increases in the average apical branch length per neuron (Fig. 2B; layer 2a neurons: p6-8 vs p12-14: p<0.0001, p12-14 vs >p30: p<0.0001; layer 2b neurons: p6-8 vs p12-14: p<0.001, p12-14 vs >p30: p<0.05; ANOVA with Holm-Sidak’s multiple comparisons test).

Layer 2b basal branches exhibited a similar pattern, the average branch length only increased significantly in the second postnatal week but not in the first postnatal week (Fig. 3B). In layer 2a neuron basal dendrites, length increase was only significant when comparing over the whole developmental period observed (Fig. 3A3; layer 2b: p6-8 vs p12-14: p<0.0001; layer 2a: p1-2 vs >p30: p0.01; ANOVA with Holm-Sidak’s multiple comparisons test).

It is obvious from Fig. 3B that the relatively small length increase in layer 2a neuron basal dendrites predominantly occurred in the second postnatal week and between postnatal weeks 2 and 5. In sum, our measurements permitted us to clearly distinguish branch addition in developmental phase one followed by elongation of individual branch segments in developmental phase two for layer 2a and layer 2b dendrites.

We identified a third developmental phase in the interval between the end of the second postnatal week (p12-14) and the fifth postnatal week (>p30). We observed a 34% reduction in the number of apical branches in layer 2b neurons and a 33% reduction in the number of stems in layer 2a neurons (Figs. 2A2 and 2A4). In layer 2b neurons, this pruning was accompanied by a halt in the increase of total dendritic branch length (Fig. 2A3) despite a significant increase in the average branch length per neuron (Fig. 2B). This resulted in a diverging developmental trajectory of the apical dendrite between layer 2a and layer 2b neurons. While the total dendritic branch length was similar until week 2, the two different developmental patterns of layer 2a and layer 2b neurons resulted in a significantly shorter apical dendritic tree in layer 2b neurons at five weeks (Fig. 2A3; layer 2a vs layer 2b at >p30: p<0.05; ANOVA with Holm-Sidak’s multiple comparisons test). Similar to their apical dendrites, layer 2b neuron basal dendrites pruned significantly after p12-14, both with respect to branch length and branch number (Fig. 3A2 and A3). Between postnatal weeks 2 and 5, we therefore defined a third developmental phase of pruning for apical and basal dendrites of layer 2b neurons and apical stems of layer 2a neurons.

To see how the three distinct developmental phases affected the spatial arrangement of branches, we plotted the distribution of branch densities as a function of distance from the soma. We observed differences between apical dendrites of layer 2a and layer 2b neurons during the first two postnatal weeks. In developmental phase 1, layer 2a neurons branched close to the soma, the distribution was single-peaked (Fig. 2C1). In contrast, layer 2b neurons also displayed a second peak of distal branching right after birth (p1-2), and during the first developmental phase determining dendritic complexity (p6-8; Fig. 2C2). During the second developmental phase of dendritic elongation, no branches were added in layer 2a and layer 2b neurons. At the same time, we observed a right shift of the peaks of apical branch density to larger distances from the soma. This indicated that length growth was not limited to dendritic tips but also affected intermediate branches. Pruning in developmental phase 3 resulted in a reduction of the second, distal peak of the layer 2b neurons, approximating the apical branch distributions of layer 2a and layer 2b neurons. The distribution of basal branch density as a function of distance from the soma was similar for both cell types, with a right shift for layer 2b neurons (Fig. 3C).

In sum, circuit-specific differences in dendritic development between layer 2a and layer 2b neurons were observed in developmental phases one (branch addition) and three (pruning). While differences in basal branch number between layer 2a and layer 2b neurons were determined in developmental phase one, development of the apical dendritic tree diverged during pruning in phase 3.

**Figure 2.**
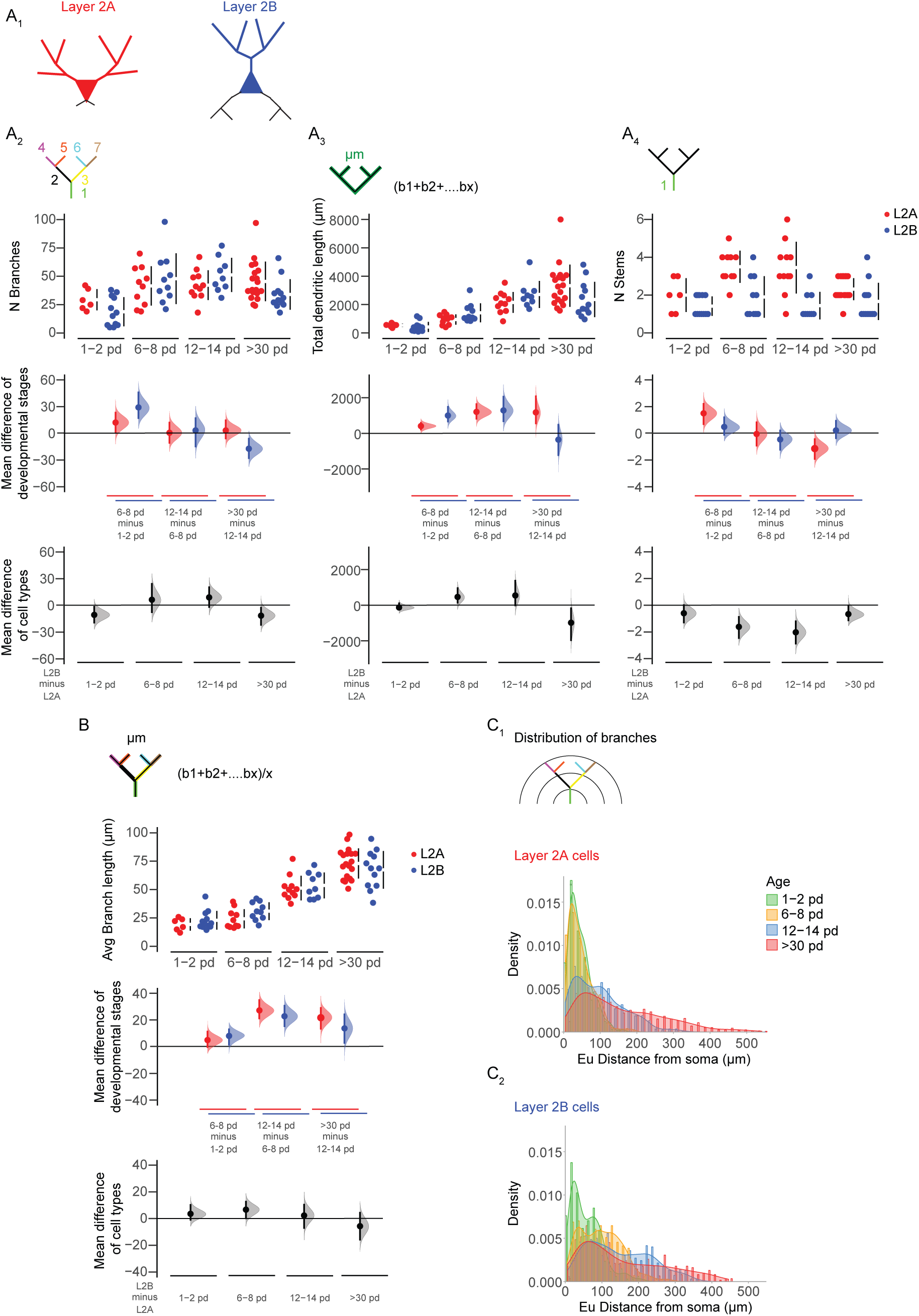
Developmental changes in the morphology of the apical dendritic tree. (A1) Visual representation of layer 2a (L2A, top, red) and layer 2b (L2B, bottom, blue) neurons. Morphological parameters are used to described growth patterns of the apical trees of these cells during development at four different ages (expressed as postnatal days, pd). Four measurements were extracted from the reconstructed cells and are displayed in Cumming estimation plots: (A2) Total number of apical branches per cell, (A3) total apical dendritic length per cell. (A4) total number of apical stems per cell and (B) Average apical branch-length per cell in µm. The raw data is plotted on the upper axes; mean differences between developmental stages are plotted on the middle axes and mean differences between the cell types are plotted on the lower axes, as a bootstrap sampling distribution. Mean differences are depicted as dots and the 95% confidence intervals are indicated by the ends of the vertical error bars. (C) Densities of the distributions of apical branches for layer 2a (C1) and layer 2b (C2) are plotted as function of the euclidean distance from the soma at four time windows: 1-2 pd (green), 6-8p pd (yellow), 12-14 pd (blue) and >30 pd (red).

**Figure 3.**
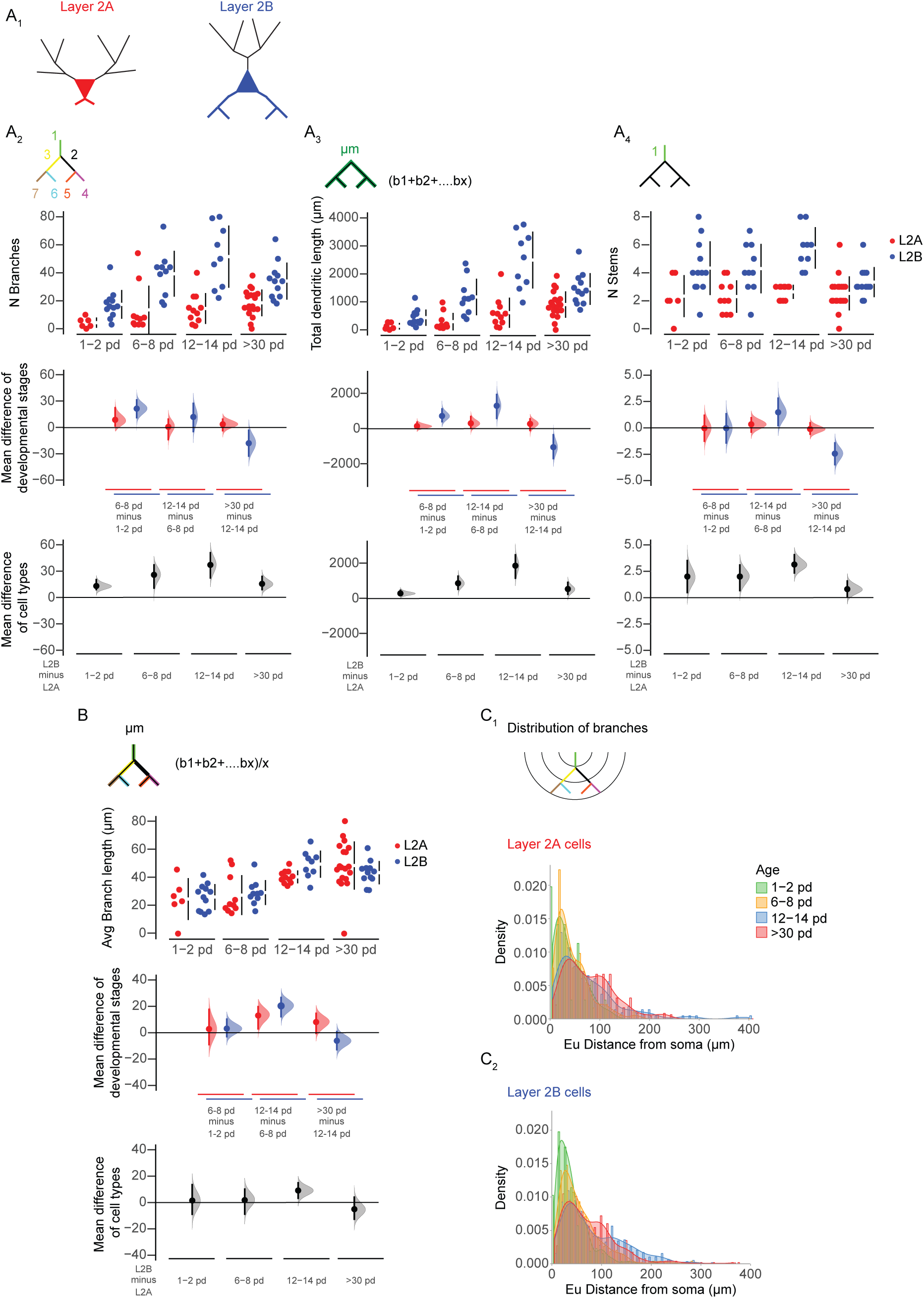
Developmental changes in the morphology of the basal dendritic tree. (A1) Visual representation of layer 2a (L2A, top, red) and layer 2b (L2B, bottom, blue) neurons. Morphological parameters are used to described growth patterns of the basal dendritic trees of layer 2a (L2A, red) and layer 2b (L2B, blue) neurons during development at four different ages (expressed as postnatal days, pd). Four measurements were extracted directly from the reconstructed cells and are shown in Cumming estimation plots: (A2) Total number of basal branches per cell. (A3) total basal dendritic length per cell in µm. (A4) total number of basal stems per cell and (B) average basal branch-length per cell in µm. The raw data is plotted on the upper axes; mean differences between developmental stages are plotted on the middle axes and mean differences between the cell types are plotted on the lower axes, as a bootstrap sampling distribution. Mean differences are depicted as dots and the 95% confidence intervals are indicated by the ends of the vertical error bars. (C) Densities of the distributions of basal branches for layer 2a (C1) and layer 2b (C2) are plotted as function of the euclidean distance from the soma at four time windows: p1-2 (green), p 6-8 (yellow), p 12-14 (blue) and >p30 (red).

### Functional connectivity during early spontaneous network activity at the mesoscale population level reflects morphological differences

The complexity of both the apical and the basal dendritic tree is determined in the first postnatal week by branch addition (developmental phase one). As dendritic structure and neuronal activity are interdependent, our next aim was to compare neuronal activity patterns during the first postnatal week between layer 2a and layer 2b neurons.

We analyzed differences between the two cell types at the mesoscale population level during immature slow spontaneous network activity patterns. Similar to the somatosensory cortex, immature slow spontaneous network activity patterns in aPCx coexist with and can be triggered by sensory inputs starting at p0 (Hoffpauir et al., 2009; Leighton and Lohmann, 2016). In acute brain slices, slow spontaneous network activity is preserved as a default state of the intrinsic recurrent network (Rigas et al., 2015).

In the juvenile circuit, layer 2a neurons are less likely to be incorporated into recurrent circuits than layer 2b neurons (Suzuki and Bekkers 2011; Wiegand et al. 2011). To date, it is unclear whether immature spontaneous network activity reflects mature connectivity patterns or acts as an unstructured global signal. Observing spontaneous network activity in the aPCx, we therefore next wanted to test whether immature spontaneous network activity early in development differentially incorporates layer 2a and layer 2b neurons. Here, we used data from Ai95-NexCre mice as this mouse line is specific for glutamatergic neurons. To assess functional connectivity between neurons, we identified all visible neurons (4755/50/39/23 neurons/fields of view/slices/mice). Based on Ca^2+^-mediated changes of the fluorescence signals, events were defined as activity-related Ca^2+^ signals distinguishable from baseline noise following the criteria stated in methods. Cells displaying events were defined as active. For neurons that were active, we extracted all events of each individual neuron. For each neuron and each event, we calculated which percentage of layer 2 neurons was coactive in the same field of view at the same time. These percentage values were then averaged over all events observed in a neuron. We plotted neuronal depth in layer 2 against the average percentage of coactive neurons in the whole layer 2 field of view. Coactivity was significantly stronger in the deep layer 2b neurons (deep third of layer 2) than in layer 2a neurons (superficial third of layer 2; Fig.4D; 1/3 layer 2 vs 3/3 layer 2: p<0.01; unpaired t-test). This was even more pronounced for the small fraction of neurons recorded in layer 1b and in layer 3 (Fig. 4C). The degree of coactivity is an indirect measure of functional connectivity. Already during the first postnatal week, we observed higher local functional connectivity in layer 2b than in layer 2a neurons. Morphologically, this scales with the more complex basal dendritic tree receiving more recurrent inputs (Haberly, 1985). We conclude that divergence of the basal dendritic tree complexity between layer 2a and layer 2b neurons is already evident in the first postnatal week and reflected by differences in functional connectivity during early spontaneous network activity.

**Figure 4.**
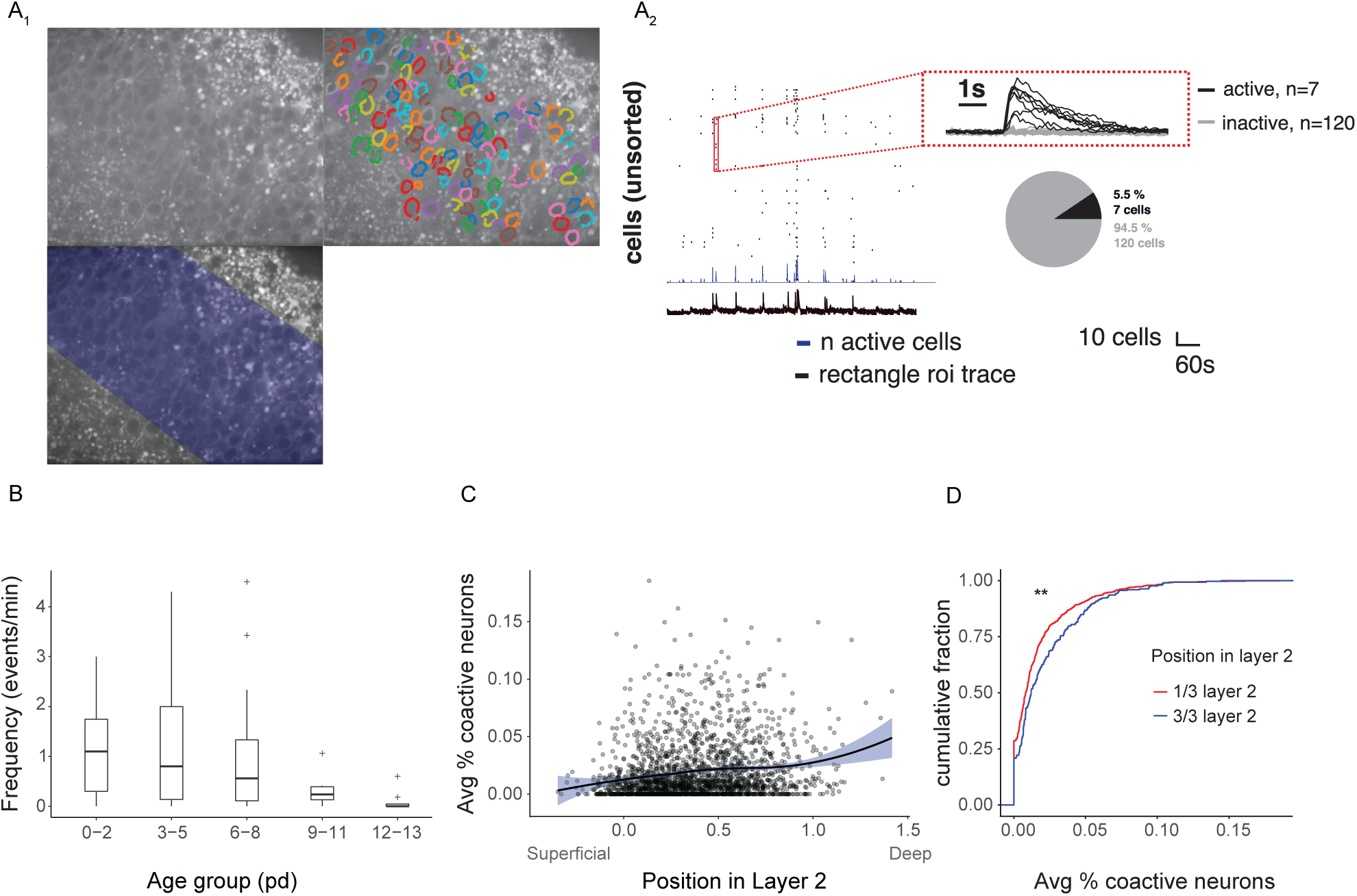
Comparison of spontaneous network activity in layer 2a and layer 2b neurons. (A1) Example of a baseline GCaMP-fluorescence image from an Ai95-NexCre mice. The field of view covers layer 2 in aPCx. Top Right: Detected cells in layer 2. Bottom: Rectangular ROI defined by the upper and lower boundaries of layer 2 for detecting global activity. (A2) Corresponding traces from the global events measured from the rectangular ROI (fluorescence: black trace, bottom) and the active cells (raster plot, blue trace for number of active cells, red inset for fluorescent traces from individual neurons in raster plot). The proportion of active versus inactive cells is indicated by pie chart for one event. (B) Frequency of spontaneous events per minute measured at 5 different age groups (expressed as postnatal days, pd; n=70/59/36 fields of view/slices/mice) (C) For each active neuron, the average percentage of coactive neurons (based on the total number of neurons in the field of view) was plotted against the position of the active neuron in layer 2 in the aPCx (data is fitted with a local polynomial regression and pooled from the first postnatal week). (D) Layer 2 was divided in 3 parts. Cumulative distribution of the average percentage of coactive neurons from (C) was plotted for the superficial third (red, 1/3 layer 2, corresponds to layer 2a neurons) and the deep third (blue, 3/3 layer 2, corresponds to layer 2b neurons). **: p<0.01.

### Pruning in layer 1a during the early critical period of sensory plasticity

Next, we wanted to further understand differences in the developmental pattern of the apical dendritic tree. Here, the most pronounced differences occurred between the end of postnatal week two and the fifth postnatal week (developmental phase 3). In this period, we observed selective pruning of layer 2b neuron apical dendrites, which did not occur in layer 2a neurons (see above).

The distinct organization of synaptic inputs to aPCx apical dendrites enabled us to relate pruning to specific circuits by grouping dendritic branches based on their position (layer 1a for branches receiving sensory inputs and layer 1b/2 for recurrent inputs). We therefore analyzed the growth and pruning patterns of apical dendrites with respect to the synaptic input layer the segments terminated in. Apical branches were categorized as branches terminating in layer 2, layer 1b (both recurrent) and 1a (sensory) for both cell types (Fig. 5A). Calretinin staining was used as a marker to delineate layer 1a (Figs. 1A3 and 4).

While basal branches of layer 2b neurons displayed clear pruning (see above), reduction of the proximal apical branches only receiving recurrent input in layers 1b and 2 did not reach statistical significance (Extended Data Figure 5-1; Kruskal-Wallis test with Dunn’s multiple comparison). Only in layer 2b neurons, distal branches that constituted sensory layer 1a circuits pruned significantly between p12-14 and >p30 (Fig. 5B1; layer 2b neurons: p12-14 vs p30: p<0.05; ANOVA with Holm-Sidak’s multiple comparisons test). Here, pruning resulted both in a significant reduction of the number of layer 1a intermediate branches and tips (Figs. 5B2 and 3; intermediate branches: layer 2b neurons p12-p14 vs > p30: p<0.05; tips: layer 2b neurons p12-p14 vs > p30: p<0.05; ANOVA with Holm-Sidak’s multiple comparisons test). Branch numbers in superficial layer 2a neurons remained stable over this developmental period (Fig. 5B). The reduction in branch number was accompanied by a significantly shorter layer 1a total dendritic length when comparing layer 2a and layer 2b neurons (Fig. 5C1; layer 2a vs layer 2b neurons at >p30: p<0.001; ANOVA with Holm-Sidak’s multiple comparisons test). Pruning was therefore limited to a specific compartment in a subpopulation of neurons.

To further understand circuit and cell-type specific pruning of the apical dendrite, we compared the average individual branch length per neuron in layer 1a of layer 2a and layer 2b neurons between postnatal weeks two and five. We observed significant branch elongation of individual layer 1a branches for both cell types (Fig. 5C2). When comparing layer 2a and layer 2b neurons, the average layer 1a branch length per neuron was similar after the second postnatal week. However, after the pruning phase, layer 1a branches in layer 2b neurons were significantly longer (Fig. 5C2; p6-8 vs. p12-14: layer 2a neurons: p<0.0001, layer 2b neurons: p<0.05; p12-14 to <p30: layer 2a neurons: p<0.01 layer 2b neurons: p<0.001; layer 2a neurons vs layer 2b neurons at >p30: p<0.05; ANOVA with Holm-Sidak’s multiple comparisons test). When analyzing the distribution change of average branch lengths during the pruning phase, we saw a shift towards longer 1a branches in both cell types, with a more pronounced shift in 2b neurons (Figs. 5D1 and D2). In layer 2a neurons, we did not observe layer 1a branch loss. Here, the length increase could be explained by branch elongation, which is a consequence of cortical growth. However, when interpreting the distribution shift in layer 2b neurons, we had to consider the pruning-related branch loss. The stronger right shift of the distribution towards longer branches in comparison to layer 2a neurons was accompanied by a significant decrease in branch number. This implied that pruning of sensory layer 1a branches in layer 2b neurons predominantly affected short branches or that surviving branches underwent enhanced length growth.

**Figure 5.**
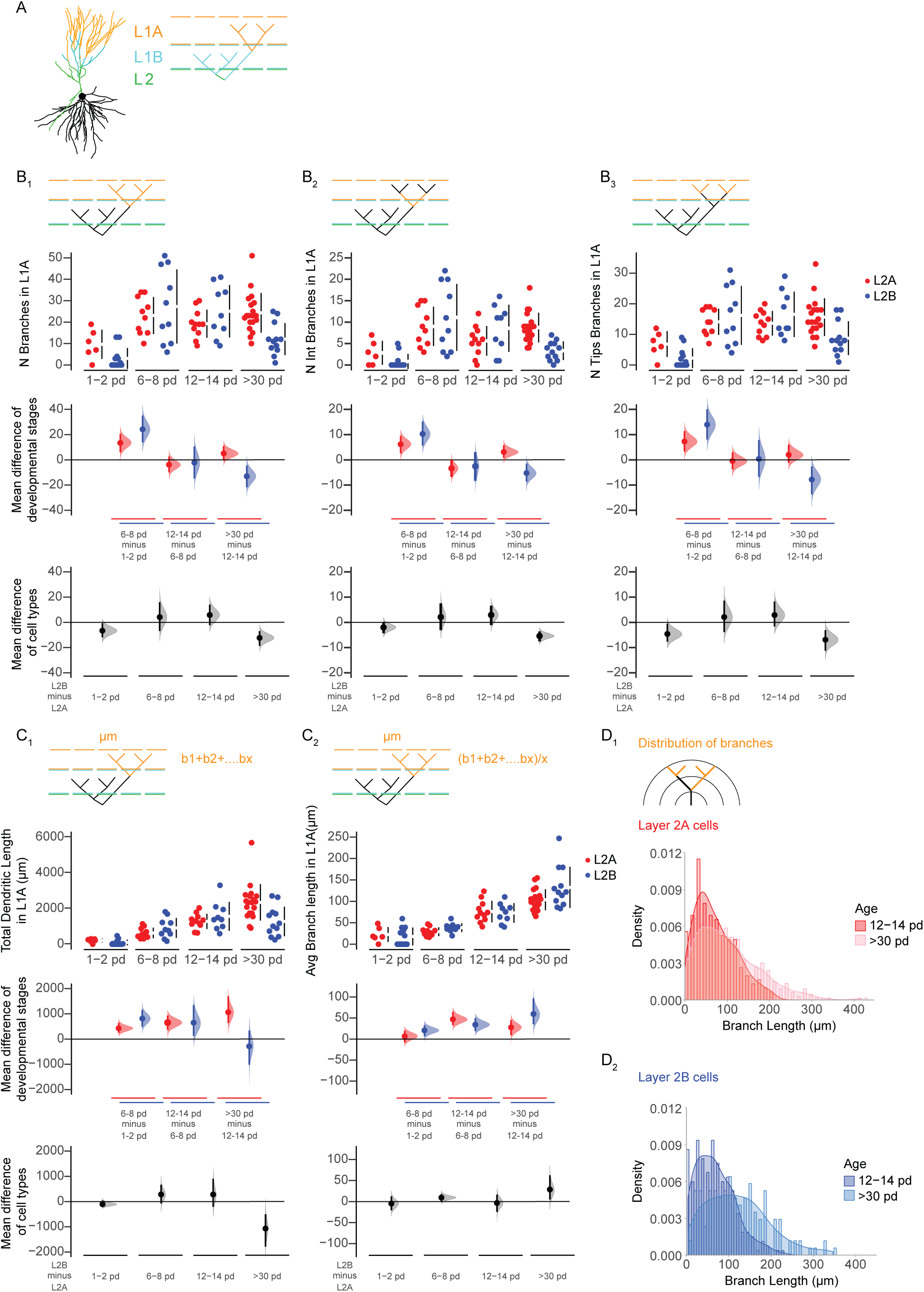
Differences in growth pattern of apical dendrites in response to layer specific synaptic inputs. (A) Example of a reconstructed cell shows the classification of the apical dendrites into three categories: branches terminating in layer 2 (L2, green), layer 1b (L1B cyan) and layer 1a (L1A, orange). (B) Growth patterns of the apical branches terminating in layer 1a are shown in Cumming estimation plots, including: Total number of branches per cell (B1), total number of intermediate branches per cell (B2), total number of tips per cell (B3), total dendritic length per cell (C1) and average branch length per cell (C2). The raw data is plotted on the upper axes; mean differences between developmental stages are plotted on the middle axes and mean differences between the cell types are plotted on the lower axes, as a bootstrap sampling distribution. Mean differences are depicted as dots and the 95% confidence intervals are indicated by the ends of the vertical error bars. (D) Densities of the distributions of the layer1a branches for layer 2a (D1) and layer 2b (D2) neurons plotted as function of branch length at two time windows: 12-14 pd (red and blue, respectively) and >30 pd (pink and light blue, respectively).

### NMDA spikes are more pronounced in layer 1a dendrites of layer 2b neurons

We next investigated the connection between the preferential pruning of short layer 1a branches and the recent discovery of supralinear dendritic integration of sensory inputs in layer 1a branches of PCx layer 2b neurons. As was recently shown in rat aPCx layer 2b neurons, clustering of synaptic inputs on the same branch as opposed to distributed input to the entire dendritic arbor significantly modified the stimulus-response behavior. Clustered (same branch) input triggered supralinear stimulus-response behavior defined as NMDAR mediated Ca^2+^ spikes (NMDA-spikes). Supralinear stimulus-response behavior resulted in large local dendritic depolarization and associated Ca^2+^ influx. Distributed inputs of similar strength evoked substantially lower levels of dendritic depolarization (Kumar et al., 2018).

We would like to relate these findings to our morphological data. Our experiments demonstrated that during postnatal development, pruning of the apical dendrites was mainly based on the loss of short layer 1a branches (Fig. 5C and D). We hypothesize that the selection bias towards longer branch segments in the sensory synaptic input space of layer 2b neurons could be a developmental mechanism supported by supralinear stimulus-response behavior. A selection bias for long branches would optimize the efficiency of the neuronal input-output function by promoting the growth and survival of branches with a higher probability of receiving clustered (same branch) input.

So far, NMDAR-dependent supralinear stimulus-response behavior has only been demonstrated in the anterior piriform cortex of rats older than four weeks (Kumar et al., 2018). Here, we hypothesize that supralinear stimulus-response behavior may play a role in the developmental branch selection observed in layer 1a dendrites of layer 2b neurons during the critical period (p12-14 to >p 30). To substantiate our hypothesis, we first wanted to test if supralinear stimulus-response behavior also occurs in in distal apical layer 1a dendrites of layer 2b neurons during postnatal week 3. We chose a developmental interval between our morphological observation points (p12-14 and >p30) as we propose that this is the time interval where supralinear stimulus-response behavior has an impact on the pruning mechanism. In contrast to layer 2b neurons, layer 2a neurons did not show pruning of short branches but equally distributed growth. If supralinear dendritic stimulus-response behavior enhanced selective pruning in distal apical dendrites of layer 2b neurons, another experimentally testable prediction would be an absence of supralinear stimulus-response behavior in layer 2a neurons.

We therefore performed combined somatic whole cell patch clamp recordings and two-photon Ca^2+^ imaging in apical layer 1a branches of layer 2a (n=5/5/5, neurons/slices/mice) and layer 2b (n=5/5/5, neurons/slices/mice) neurons. Synaptic stimulation was achieved by focal electrical stimulation with theta-glass electrodes (Figs. 6 A4 and B4). Using Ca^2+^ imaging, we first identified a focal stimulation spot. Stimulation strength was then linearly increased. When plotting the area under the curve (AUC) of the EPSP in layer 2b neurons, we observed a distinct supralinear increase at specific stimulation strengths, which was abolished after washing in of the NMDAR-antagonist APV (Fig. 6A3). In addition, this increase displayed the typical shape of an NMDA-spike (Fig. 6A1). We performed Ca^2+^ imaging in parallel and could further observe that the nonlinear enhancement of the EPSP AUC was accompanied by a branch-specific increase in spatial spread and amplitude of the Ca^2+^ transient (Fig. 6A5). These hallmarks of supralinear dendritic NMDA spikes were not observed in layer 2a neurons (Fig. 6B). Next, we plotted the AUC and the amplitude of layer 2a and layer 2b neuron EPSPs before and after wash-in of APV. We observed a significant decrease of both parameters in layer 2b neurons. In contrast, NMDAR-block by APV did not affect AUCs and amplitudes of EPSPs in layer 2a neurons (Figs. 6 C1and C2; layer 2b, AUC: pre vs. post: p<0.05; layer 2a, pre vs. post: p=0.18; amplitude: layer 2a, pre vs. post: p<0.05; layer 2b, pre vs. post: p=0.7; paired t test). At the stimulation site, the stimulation-evoked Ca^2+^ signal of both layer 2a and layer 2b neurons was significantly reduced by APV (Fig. 6D; layer 2b, pre vs. post: p<0.01; layer 2a, pre vs. post: p<0.05; ratio paired t test). Under NMDAR-block by APV, the AUC and the amplitude of the EPSP constitute a readout of the AMPA-type glutamate receptor mediated depolarization. Consequently, the absolute value of the AUC and the amplitude in APV reflected the synaptic input strength of our stimulation. These values were not significantly different between layer 2a and layer 2b neurons in our sample, indicating comparable stimulation strength for both cell types (Fig. 6C; AUC: layer 2a post vs. layer 2b post: p=0.92; amplitude: layer 2a post vs. layer 2b post: p=0.25, t-test).

We conclude that, in contrast to layer 2b neurons, layer 2a neurons do not display supralinear stimulus-response behavior in layer 1a at comparable input strengths. In layer 2b neurons, supralinear stimulus-response behavior can be observed during the developmental period of dendritic pruning of short dendritic segments. In layer 2a neurons, we neither observe pruning nor supralinear stimulus-response behavior in the same developmental period.

**Figure 6.**
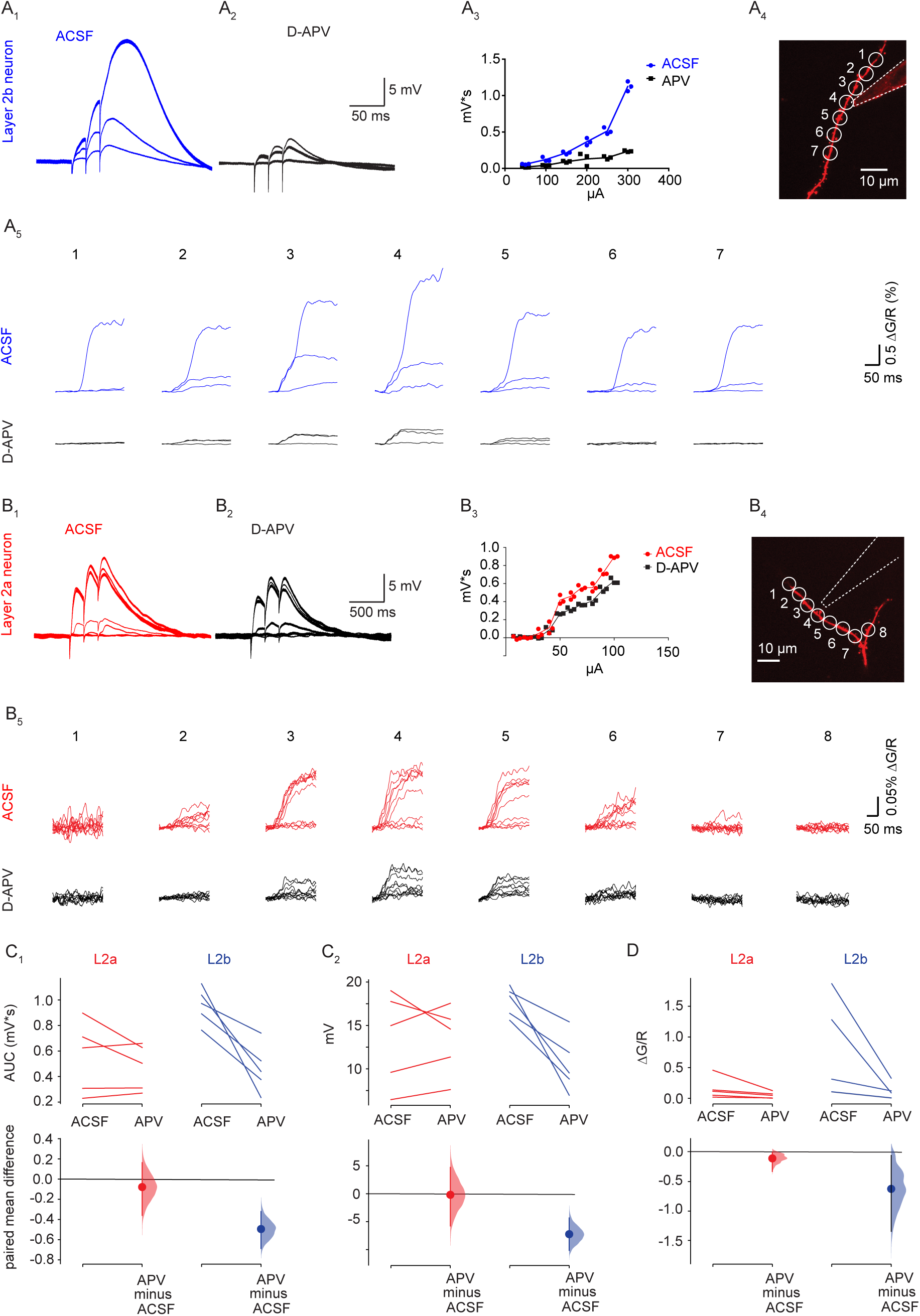
Dendritic NMDA-spikes can only be observed in layer 2b neurons. (A) displays a representative layer 2b neuron, (B) a representative layer 2a neuron. (A1/B1) Overlay of electric responses measured at the soma upon electrical stimulation with linear strength increase in layer 1a. (A2/B2) Response to same stimulus after application of APV. (A3/B3) Plot of stimulation strength against area under the curve from the same cell. (A4/B4) Imaged dendritic branch and position of the stimulation electrode. (A5/B5) Fluorescence traces of Ca2+ responses at ROIs outlined in A4/B4 under baseline conditions (upper rows, blue and red) and after application of APV (lower rows, black). Increasing stimulation strengths are overlayed. (C) and (D) Cell-type specific changes to NMDAR-block with APV are plotted as Cumming estimation plots. The raw data is plotted on the upper axes, each pair is connected by a line; each mean difference is plotted on the lower axes as a bootstrap sampling distribution. Mean differences are depicted as dots and the 95% confidence intervals are indicated by the ends of the vertical error bars. (C1) Area under the curve (AUC) before and after application of APV in layer 2a neurons (red) and layer 2b neurons (blue). (C2) EPSP amplitude before and after application of APV in layer 2a neurons (red) and layer 2b neurons (blue). (D) Fluorescent Ca2+ response before and after application of APV in layer 2a neurons (red) and layer 2b neurons (blue). Please note that the we measured Ca^2+^ responses in 5 layer 2b neurons, however the datapoints of 2 measurements were very close and occlude each other.

We next applied computational modeling to test if the probability of supralinear integration by clustered inputs could indeed scale with branch length over the range of branch lengths measured in this study. As a quantitative framework, we used the range of average branch lengths we observed at the beginning of the critical period (p12-14; 40 to 110 µm). The average total dendritic branch length (1800 µm) was set constant for all model neurons. We estimated the layer 1a input density based on a recent comprehensive quantitative description of mouse piriform cortex (Srinivasan and Stevens, 2018). Based on Srinivasan and Stevens, we extrapolated that the whole population of 3700 bulbar glomeruli makes 2366 synapses with each individual neuron. Consequently, one glomerulus makes on average 0.64 synapses with each neuron. 109 coincidently activated glomeruli would therefore activate 70 synapses on a layer 2b neuron, which we defined as the upper limit of coincident inputs. Fig. 7B displays the results of our calculation. We plotted the probability of dendritic NMDA-spikes evoked by clustering of more than 10 synapses on an individual branch as a function of the average branch length per neuron. This was repeated for different numbers of coincident synaptic inputs (see methods for details). The dendritic spike probability increased with increasing branch length. Within our parameter space, clustering probability and the resulting dendritic spiking increased from close to 0 to up to 17% when comparing the shortest and longest average branch length we observed in our dataset. Quantitatively, our results suggest that, during our developmental time window, dendritic NMDA spikes are more likely to occur in long dendritic segments than short ones. It is therefore plausible that the supralinear stimulus-response behavior observed exclusively in layer 2b neurons could constitute a selection mechanism optimizing the efficiency of the neuronal input-output function. This would occur by promoting the growth and survival of branches with a higher probability of receiving clustered (same branch) input.

**Figure 7.**
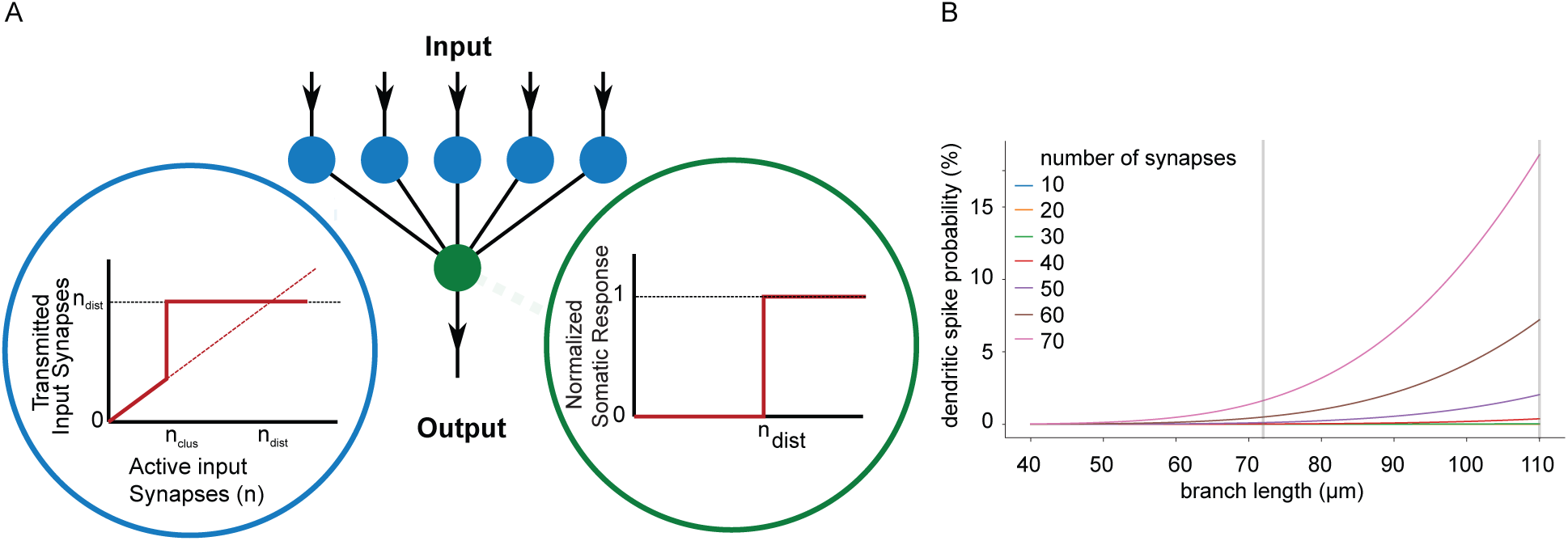
(A) Schematic neuron model as two-layered neural network, consisting of a dendritic activation layer (blue circles) and a somatic activation layer (green circles). The activation functions for the respective layers are shown in the insets. The activation of the soma is modeled as simple step function (if the input surpasses a certain threshold, the soma responds). The activation of every separate dendrite branch is modeled as an overlay of a linear response and a (non-linear) step function, to mimic the excitability of dendritic branches. If a dendritic branch is excited (due to a large enough number of same branch active input synapses) it transmits, triggering a somatic response. (B) The response probability increases as function of the branch length, an effect that becomes more apparent for different increasing numbers of active input synapses S.

### Discussion

Here, we performed a morphometric and functional analysis of postnatal dendritic development in aPCx sensory and recurrent circuits. We compared developmental patterns of excitatory neurons in the sublayers 2a and 2b. The two neighboring sublayers differ with respect to sensory and recurrent wiring. We could identify a timeline defined by three developmental phases: 1. Branch addition (developmental phase one, postnatal week one). 2. Branch elongation (developmental phase two, postnatal week two). 3. Branch pruning (developmental phase 3, postnatal weeks 3-5). We discovered circuit and sublayer-specific differences in dendritic development in developmental phases one and three. In developmental phase one, layer 2a neuron basal dendrites incorporated in recurrent circuits branched significantly less than layer 2b neuron basal dendrites. This was accompanied by lower functional connectivity of layer 2a neurons during spontaneous immature network activity. In developmental phase three, pruning of apical dendrites receiving layer 1a sensory inputs was only observed in layer 2b neurons, but not in layer 2a neurons. Pruning was clearly biased towards shorter dendritic branches. Using electrophysiology, Ca^2+^ imaging and modeling, we demonstrated how NMDAR-dependent supralinear stimulus-response behavior during phase 3 could support the survival and growth of long dendritic branches. Nonlinear dendritic properties could therefore be involved in dendritic development.

#### Distinction between layer 2a and layer 2b neurons

Distinguishing principle cell types in layer 2 poses a ‘lumping versus splitting’ problem. Our hard segregation into layer 2a and layer 2b neurons based on location on the vertical axis qualifies as a ‘lumping’ approach. Along the vertical axis of layer 2, we can distinguish at least two different types of principal cells based on their position, connectivity and morphology. Based on this distinction, principal neurons in layer 2 have been hypothesized to represent two parallel streams of olfactory information processing. Superficial neurons receive predominantly sensory inputs, whereas deep neurons receive both sensory and recurrent inputs (Suzuki and Bekkers 2011; Wiegand et al. 2011).

When segregating layers 2a and 2b as done here, this dichotomy is complicated by the observation that there seems to be a vertical gradient between so-called semilunar cells with no basal dendrites and superficial pyramidal cells with elaborate basal dendrites (Bekkers and Suzuki 2013). Our detailed morphological reconstructions and analysis support the idea of a continuum between these differentially wired types of neurons. We observed short or absent basal dendrites in layer 2a and increasing basal dendritic complexity in layer 2b. It is established that basal dendrites primarily receive local recurrent connections (Haberly, 1985; Luskin and Price, 1983). Therefore, basal dendritic complexity is a robust criterion for distinguishing between neuronal subtypes receiving different amounts of recurrent input. We were therefore able to differentiate between neurons in layer 2a and layer 2b on a vertical axis. This is important, as the intrinsic properties we observed along the vertical axis of layer 2 were more homogenous, which is different from previously published results in mice (Suzuki and Bekkers, 2011, 2006) and rats (Wiegand et al., 2011). Of all our age groups, our p12-14 data most closely matches the p13-p30 observation window used for the initial description of electrophysiological differences between superficial and deep layer 2 neurons in a mouse strain closely related to ours (Suzuki and Bekkers, 2006). While we found a similar trend regarding the difference in input resistance and AP threshold, our cell population segregated less clearly regarding the initial firing frequency. In addition, superficial neurons in our dataset were as hyperpolarized as deep neurons, whereas more depolarized superficial layer 2 neurons were recorded in earlier studies (Table 1, Extended Data Table 1-1). We can list a number of reasons that could underly the observed differences: Differences in sampling (covering the whole extent of layer 2 in our case rather than focusing on the lower layer 2/3 and upper border as has been done previously, a more stringent definition of the deep layer 2 border that minimizes layer 3 neurons displaying more pronounced bursting); different positions on the anterior-posterior axis; holding potential (as opposed to earlier studies, our layer 2b neurons were characterized at -60mV, where the burst-mediating T-Type Ca^2+^ channels are most likely inactivated (Joksimovic et al., 2017)); slicing angle and changes in intrinsic electrical properties when using a KMESO4-based intracellular solution as opposed to a KGLUC-based intracellular solution in our case (Kaczorowski et al., 2007).

Recent studies have extended the parameter space for splitting aPCx layer 2 neurons: long-range tracing studies identified layer-specific differences in axonal projection patterns (Chen et al., 2014; Diodato et al., 2016; Mazo et al., 2017). Different excitatory cell types have also been identified at the level of genetic markers. Genetic marker expression profiles of superficial neurons constituting layer 2a further support the ‘hard’ location-based segregation principle applied here, as, based on published representative examples, they clearly seem to delineate a layer2a/layer2b border (Bolding et al., 2019; Choy et al., 2017; Diodato et al., 2016).

#### Critical period imprinting and pruning of layer 2b neuron dendrites in the non-topographic aPCx

The aPCx is a sensory brain region with a simple and evolutionary conserved structure that predates the development of the sensory neocortex (Bekkers and Suzuki, 2013). A central difference between sensory neocortex (auditory, visual and somatosensory) and the aPCx is the spatial organization of afferent input. Sensory neocortex is a topographic circuit: nearby peripheral neurons share similar and predictable representations of sensory space and contact neighboring cortical neurons. These neighboring neurons have a high degree of local connectivity. In contrast, the PCx is a distributed, non-topographic circuit, where both the representation of sensory space and the recurrent connectivity are unpredictable and dispersed (Srinivasan and Stevens, 2018). This diffuse organization of the palaeocortical PCx is likely to reflect the primordial structure of cerebral cortices in reptiles prior to the evolution of isocortex in synapsids and later mammals (Fournier et al., 2015). These differences in the afferent input structure could affect dendritic growth patterns in palaeo- and neocortex. In the following, we would therefore like to compare our data with published neocortical patterns of dendritic growth.

The determination of branch complexity during the first postnatal week corresponds to dendritic growth patterns identified in layer 2/3 and layer V pyramidal neurons in rodent neocortex (Maravall, 2004; Petit et al., 1988; Romand et al., 2011). The determination of branch complexity in the first postnatal week can therefore be considered a common design principle of neocortex and palaeocortex.

As in our palaeocortical dataset, length increase of neocortical dendrites was significant between the first and second postnatal week both in basal and apical dendrites. In the neocortex, there was no further increase in the period until p30 (Petit et al., 1988; Romand et al., 2011). The developmental trajectories of neocortex and palaeocortex diverge in developmental phase three during the critical period of sensory plasticity in the aPCx (between p12-14 and > p30): In the palaeocortical aPCx, we observed a reduction in total dendric length and in dendritic branch number in apical and basal dendrites beween p12-14 and p30. In neocortical layer 5 neurons, a decrease in branch number was observed earlier, before the onset of the critical period between p7 and p14 in apical but not in basal dendrites and was attributed to filopodial, not dendritic pruning (Romand et al., 2011).

We propose that an overabundance of dendrites at the beginning of the critical period in aPCx could be a consequence of the requirements of a non-topographic afferent input structure. Overabundance of distal apical layer 1a dendrites increases the combinatorial space for different glomerular input combinations. A surplus of basal dendrites results in more recurrent connections. We would have a larger probability of generating circuit motifs of recurrently connected neurons sharing similar sensory inputs. The enlarged combinatorial space maybe necessary in a distributed circuit like the PCx. In contrast, the topographic neocortex displays local clustering of similar inputs. This will result in a higher probability of the clustering of similar inputs on interconnected neurons. An effect of neocortical topographical organisation may therefore be the effective wiring of significant features of sensory space. This would obviate the need for the metabolically expensive generation of superfluous dendritic branches during neocortical development. In the palaeocortical aPCx, the overabundance of dendrites combined with pruning could therefore enhance the initial combinatorial space and compensate for the lack of effective wiring by topography biases. Interestingly, the time window of pruning in layer 1a matches the critical period of NMDAR-dependent circuit specific sensory synaptic plasticity in layer 1a (Kevin M Franks and Isaacson, 2005; Poo and Isaacson, 2007). We therefore propose that in layer 2b neurons, the critical period in the aPCx is accompanied by mediated circuit specific pruning and remodeling of the distal apical dendritic tree receiving sensory inputs in layer 1a. In the future, it will be interesting to test if initial dendritic overabundance followed by pruning is a general feature of non-topographically ordered cortices.

#### Possible contribution of supralinear dendritic integration to pruning

In layer 2b neurons, the enlargement of the olfactory coding space at the beginning of the critical period is followed by a reduction of dendritic branches. Here, we hypothesize that dendritic NMDAR-dependent supralinear stimulus-response behavior could serve as an underlying mechanism for this pruning process. Our rationale is that assuming constant synaptic density in layer 1a, longer dendritic branches will result in a higher probability for regenerative NMDA-spikes evoked by clustered inputs. This is backed by a recent publication demonstrating that inputs clustered on the same layer 1a branch in aPCx result in supralinear integration irrespective of the distance between the inputs. In addition, large NMDAR-mediated Ca^2+^ signals occur during NMDA-spikes (Kumar et al., 2018). NMDAR-mediated Ca^2+^ signals are generally considered to promote dendritic growth during development (Konur and Ghosh, 2005). Our data therefore is compatible with an NMDA-spike dependent selection and optimization process for apical dendritic layer 1a branches of layer 2b neurons. NMDA-spikes occur with a higher probability on longer branches, the related Ca^2+^ rise would serve as a dendritotrophic signal. Such a mechanism could promote the observed survival and elongation of long branches at the expense of short branches. The result would be the structural self-amplification of dendritic NMDA-spikes by promoting branches with a higher probability for clustered inputs.

Although our Ca^2+^ imaging data in layer 1a strongly suggests local dendritic NMDA-spikes in layer 1a, our experiments do not exclude a significant contribution of recurrent inputs to the enlarged EPSP amplitudes we interpret as dendritic NMDA-spikes. With respect to pattern completion in the aPCx, such a mechanism is plausible. Supralinear stimulus response behavior of layer 1a input evoked by coincident sensory and recurrent inputs recruited by our extracellular stimulation would also generate large local Ca^2+^ transients in the activated dendritic branches. This would both be the case when the inputs share the same (long) branch and project onto different branches. It is therefore also possible that supralinear stimulus response behavior could be based on coincident activity between distal and proximal dendritic compartments. The supralinear integration of sensory and recurrent inputs could serve as a plasticity inducing coincidence detection mechanism. This mechanism would be comparable to the plateau potential evoked by coincident proximal and distal inputs in hippocampal CA1 pyramidal neurons (Takahashi and Magee, 2009). In the aPCx, such a mechanism would amplify synaptic connections between ensembles of recurrently connected neurons that share similar sensory inputs. Interestingly, we also observed an overabundance of the recurrently connected basal dendrites in layer 2b neurons. These basal dendrites underwent pruning together with layer 1a dendrites. It is tempting to speculate that pruning skips apical and basal branches of interconnected layer 2b neurons sharing similar sensory inputs. This could optimize the input-output function of dendritic branches coding for relevant features in interconnected neurons sharing sensory and recurrent inputs. Ideally, longitudinal in vivo studies of dendritic Ca^2+^ dynamics and the related dendritic growth during the critical period would be needed to further support this hypothesis.

In contrast, layer 2a neuron apical dendrites in layer 1a do neither display supralinear stimulus response behavior of layer 1a inputs nor pruning. In addition, they have a larger dendritic tree in the sensory layer 1a. This fits well with the stronger incorporation into sensory circuits these neurons display (Suzuki and Bekkers, 2011; Wiegand et al., 2011) and may result in larger olfactory receptive fields that are developmentally hardwired.

In sum, we provide evidence for circuit-specific mechanisms of dendritic development in the aPCx. We demonstrated that different developmental trajectories of the dendritic tree in layer 2a and layer 2b neurons relate to differences in circuit incorporation. Our data therefore support the concept that structural and functional differences between layer 2a and layer 2b neuron dendritic trees determine their distinct functions in the aPCx.

## Acknowledgements

We would like to thank Robert Sachdev and Matthew Larkum for the Ai95 mouse line. We would further like to thank Anke Schönherr and Susanne Rieckmann for excellent technical assistance.

## Extended Data

**Extended Data Figure 1-1.**
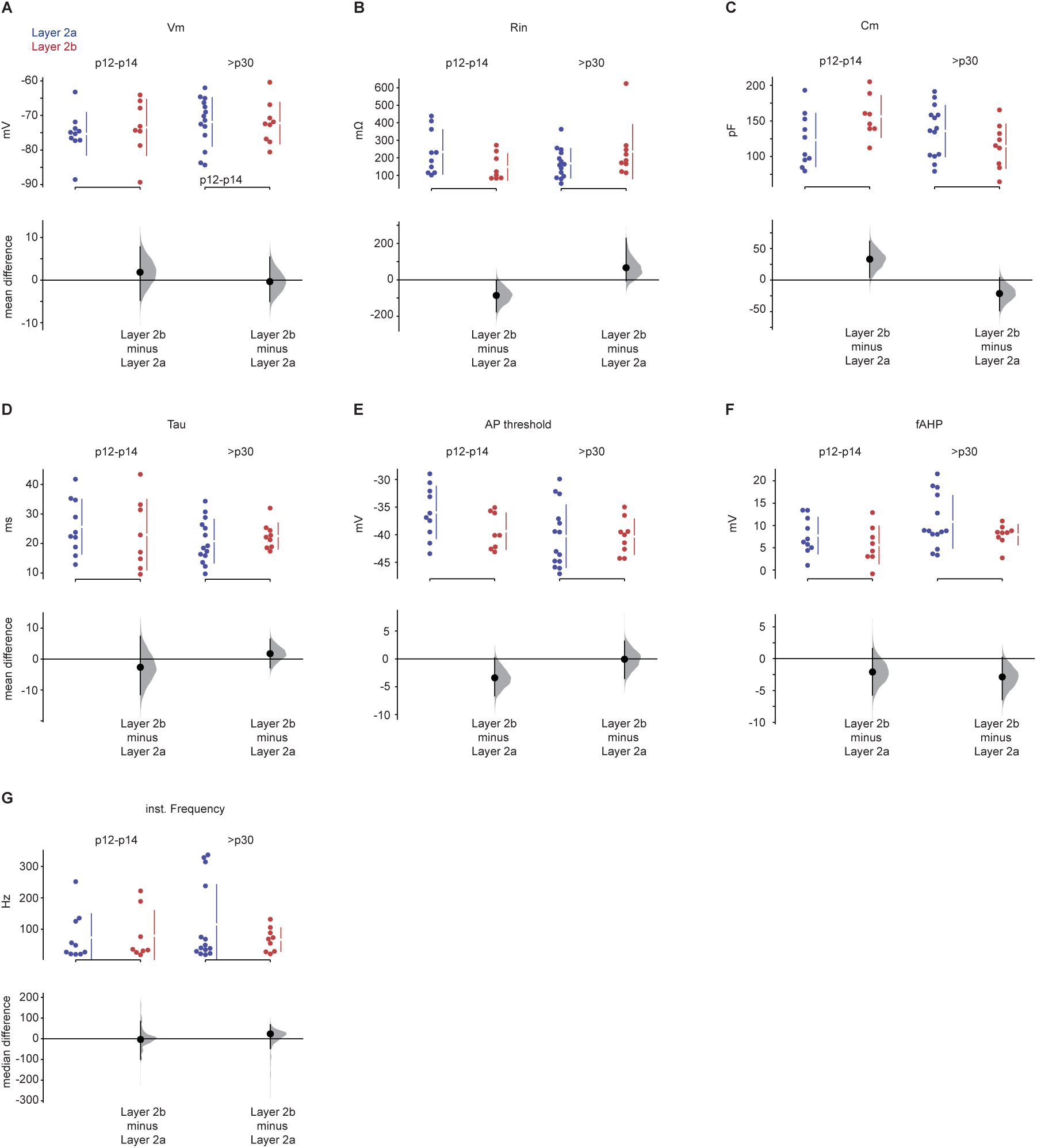
Statistical analysis of the correlation between the total basal dendritic length and the vertical position of the cells in layer 2 at four time windows: 1-2 pd, 6-8 pd, 12-14 pd and >30 pd.

**Extended Data Table 1-1.** A to F: The mean differences of intrinsic electrophysiological parameters of layer 2a (blue) and layer 2b (red) neurons at p12-14 (left) and at >p30 (right) are shown in Cumming estimation plots. The raw data is plotted on the upper axes; each mean difference is plotted on the lower axes as a bootstrap sampling distribution. Mean differences are depicted as dots and the 95% confidence intervals are indicated by the ends of the vertical error bars. (A) refers to resting membrane potential Vm, (B) to the input resistance Rin, (C) to the membrane capacitance Cm, (D) to the membrane time constant tau, (E) to the AP threshold and (F) to the fast afterhyperpolarizing potential fAHP. (G) The median differences of the instantaneous firing frequency of layer 2a (blue) and layer 2b (red) neurons at p12-14 (left) and at >p30 (right) are shown in Cumming estimation plots. The raw data is plotted on the upper axes; each median difference is plotted on the lower axes as a bootstrap sampling distribution. Median differences are depicted as dots and the 95% confidence intervals are indicated by the ends of the vertical error bars.

| Data | Type of Test | Comparison | r | P-value | Significant |
| --- | --- | --- | --- | --- | --- |
| Figure 1C1 | Spearman r | Location L2 vs Basal Length P1-2 | 0.6161 | 0.0065 | ** |
| Figure 1C2 | Spearman r | Location L2 vs Basal Length P6-8 | 0.6793 | 0.001 | *** |
| Figure 1C3 | Spearman r | Location L2 vs Basal Length P12-14 | 0.8358 | < 0.0001 | **** |
| Figure 1C4 | Spearman r | Location L2 vs Basal Length P>30 | 0.5442 | 0.0016 | ** |

**Extended Data Table 1-2.**
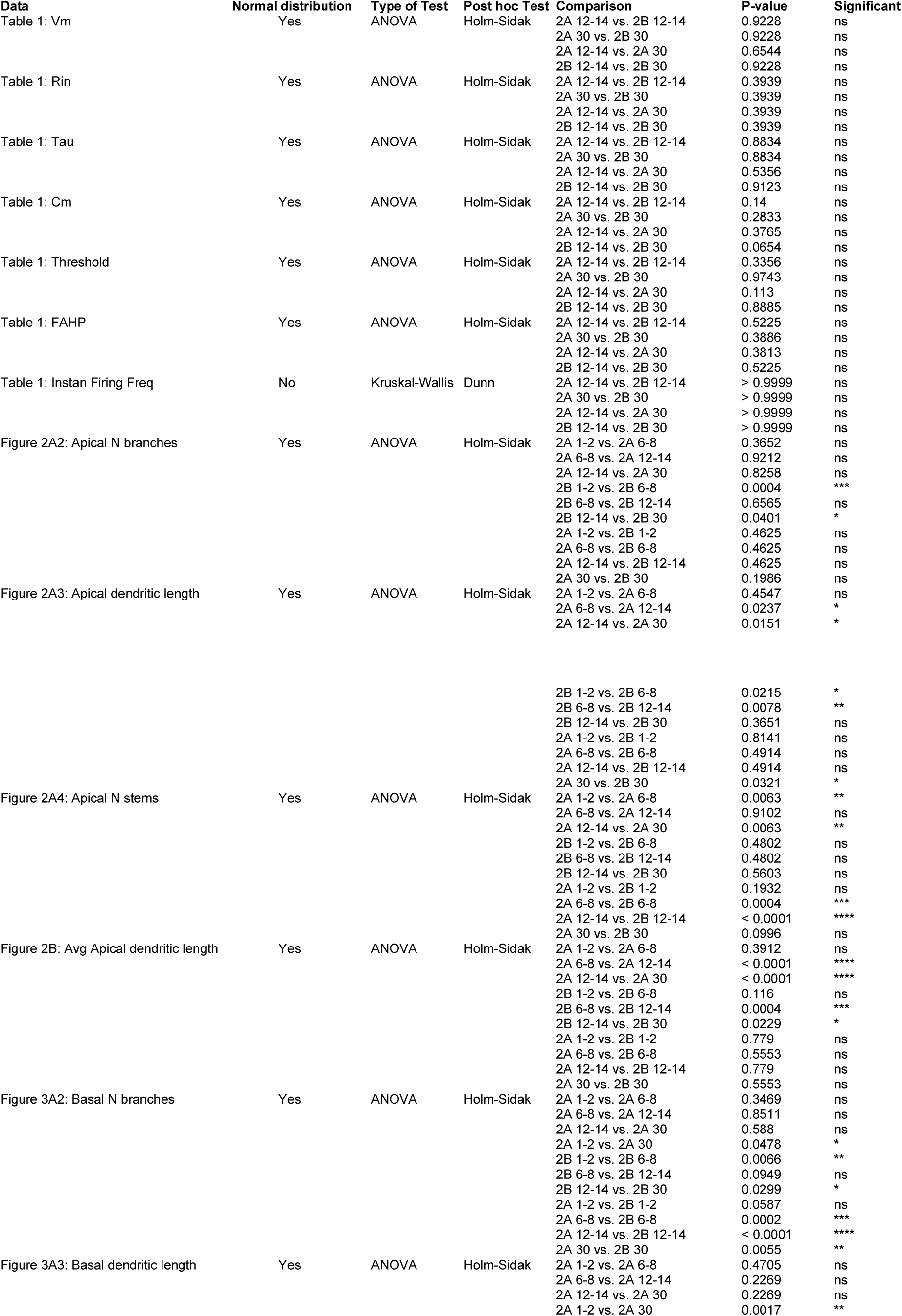
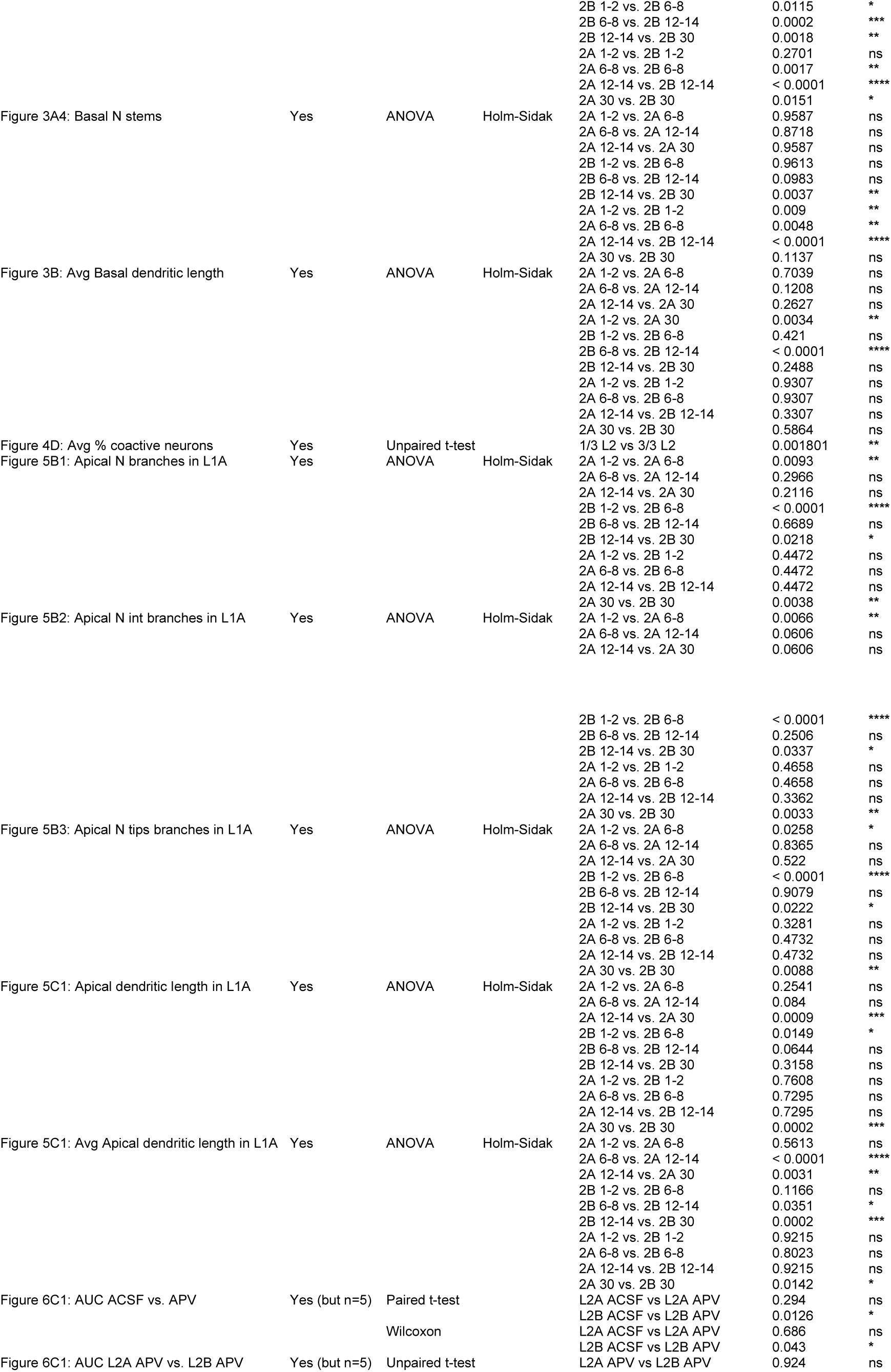
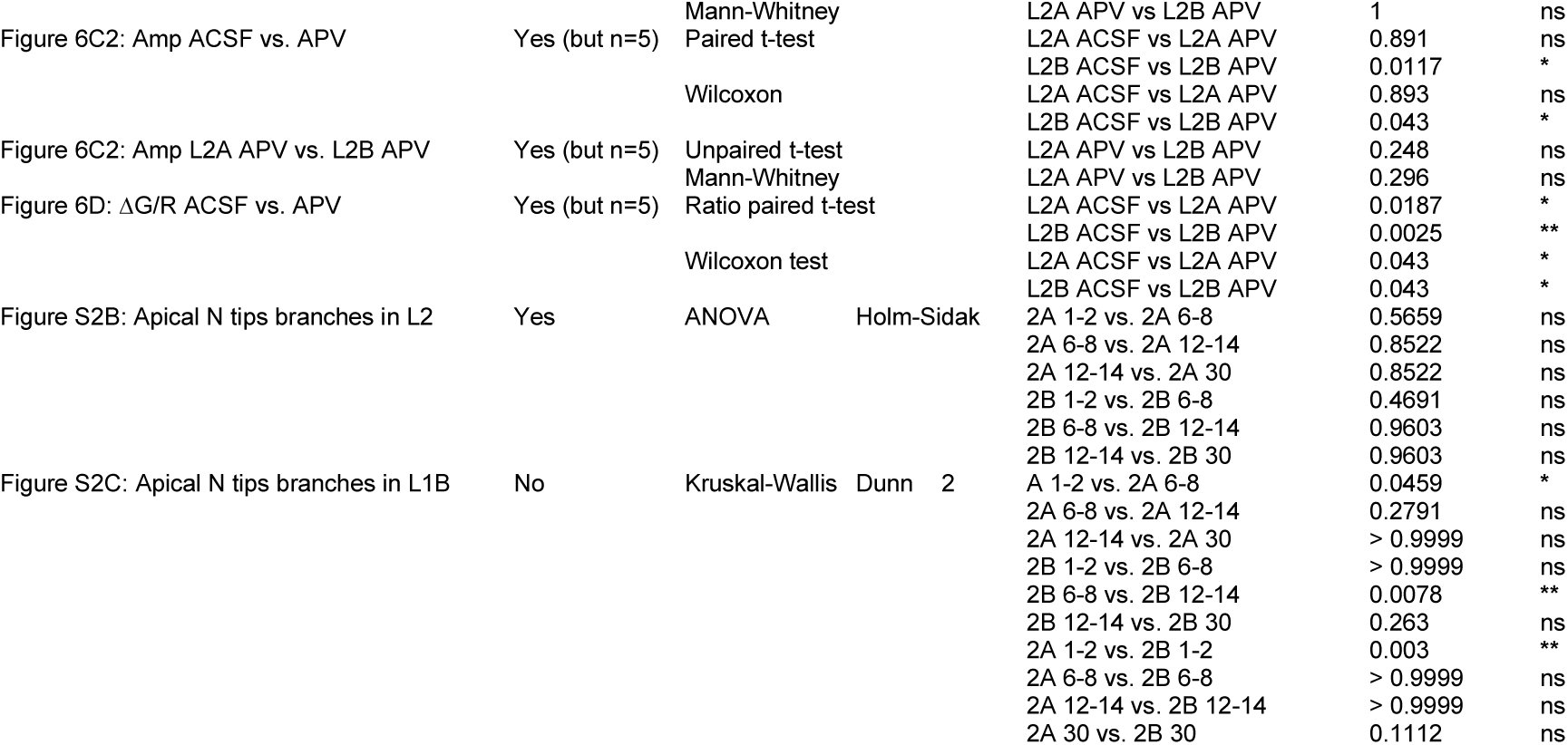
Statistical analysis of the intrinsic electrical properties and morphological parameters of layer 2a and layer 2b neurons at p12-14 and > p30.

**Extended Data Figure 5-1.**
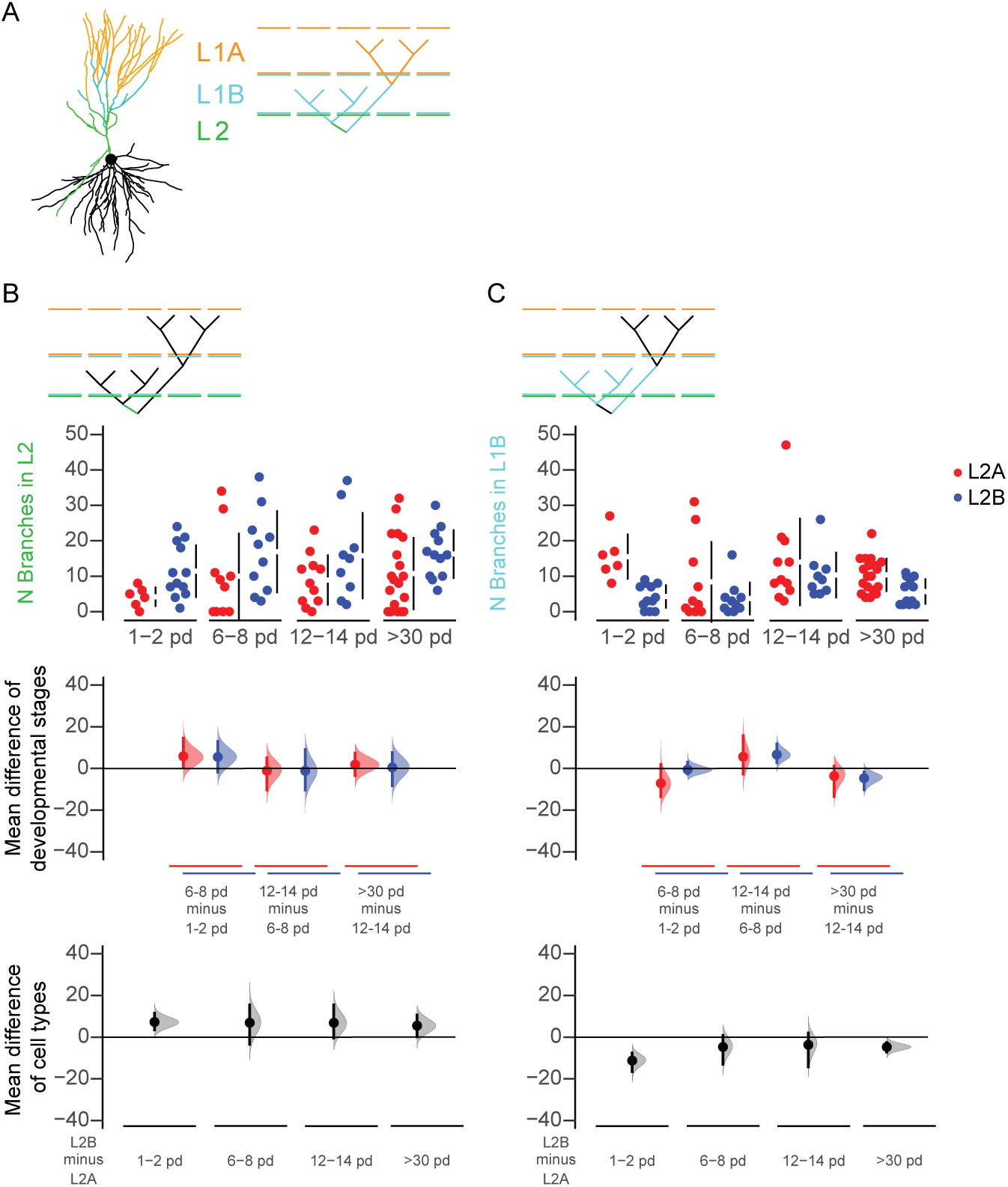
Layer specific terminating branches. (A) Example of a reconstructed cell shows the classification of the apical dendrites into three categories: branches terminating in layer 2 (L2, green), layer 1b (L1B cyan) and layer 1a (L1A, orange). (B-C) The total number of branches terminating in layer 2 (B) and layer 1b (C) for layer 2a (L2A, red) and layer 2b (L2B, blue) neurons are shown in Cumming estimation plots. The raw data is plotted on the upper axes; mean differences between developmental stages are plotted on the middle axes and mean differences between the cell types are plotted on the lower axes, as a bootstrap sampling distribution. Mean differences are depicted as dots and the 95% confidence intervals are indicated by the ends of the vertical error bars.

